# Development of a Ribosome Profiling Protocol to Study Translation in the yeast *Kluyveromyces marxianus*

**DOI:** 10.1101/2022.02.06.478964

**Authors:** Darren A Fenton, Stephen J Kiniry, Martina M Yordanova, Pavel V Baranov, John P Morrissey

**Author notes:** Corresponding author Prof John Morrissey, School of Microbiology, University College Cork, Cork T12 K8AF, Ireland **email:** |.

## Abstract

*Kluyveromyces marxianus* is an interesting and important yeast because of particular traits like thermotolerance and rapid growth, and applications in food and industrial biotechnology. Knowing how *K. marxianus* responds and adapts to changing environments is important to achieve a full understanding of the its biology and to develop bioprocesses. For this, a full suite of omics tools to measure and compare global patterns of gene expression and protein synthesis is needed. Whereas transcriptome analysis by RNA-Seq quantifies mRNA abundance, ribosome profiling allows codon-resolution of translation on a genome-wide scale by deep sequencing of ribosome locations on mRNAs and is emerging as a valuable tool to study translation control of gene expression. We report here the development of a ribosome profiling method for *K. marxianus* and we make the procedure available as a step by step protocol. To aid in the analysis and sharing of ribosome profiling data, we also added the *K. marxianus* genome as well as transcriptome and ribosome profiling data to the publicly accessible GWIPS-viz and Trips-Viz browsers. Users are able to upload custom ribosome profiling and RNA-Seq data to both browsers, therefore allowing easy analysis and sharing of data. As many studies only focus on the use of RNA-Seq to study *K. marxianus* in different environments, the availability of ribosome profiling is a powerful addition to the *K. marxianus* toolbox.

**Graphical abstract:** Development of a Ribosome Profiling protocol to study gene expression in the thermotolerant yeast *Kluyveromyces marxianus*.

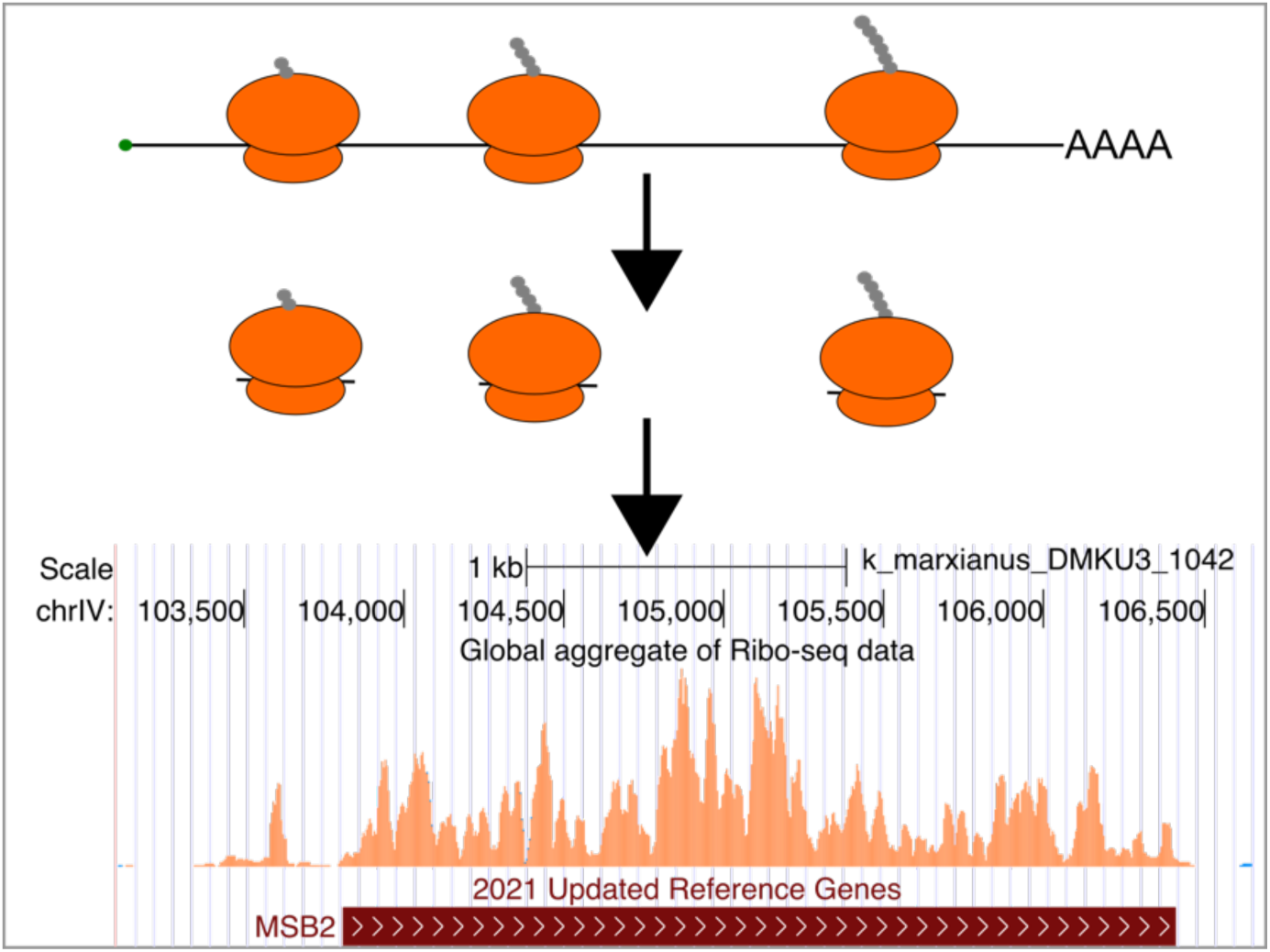

## Introduction

As with other microbes, yeasts have evolved elaborate mechanisms to sense and respond to changing extracellular and intracellular environments. External influences include phenomena such as altered nutrient availability, toxic molecules, temperature fluctuations, low pH and high osmotic pressure, while internally, cells can experience changes such as reduced intracellular pH, ion fluxes, energy depletion or nutrient starvation (Martínez-Montañés, Pascual-Ahuir and Proft 2010; Broach 2012; Ljungdahl and Daignan-Fornier 2012; Morano, Grant and Moye-Rowley 2012; de la Torre-Ruiz, Pujol and Sundaran 2015; Sui *et al*. 2015; Taymaz-Nikerel, Cankorur-Cetinkaya and Kirdar 2016). The best-studied response mechanisms in yeast involve sensor systems, signal transduction pathways, and changes in gene expression (de Nadal and Posas 2010). Ultimately, this gives rise to a new set of proteins that enable the cell to adapt, if necessary, and to survive as well as proliferate in this new environment. Dissecting these response mechanisms is central to understanding the fundamental biology of a species, but it is also important for the development of yeast for biotechnological applications (Liu and Nielsen 2019). Yeasts are used for diverse applications in the food, biopharma and industrial biotechnology sectors (Arevalo-Villena *et al*. 2017; Nandy and Srivastava 2018; Parapouli *et al*. 2020) and, very often, they need to perform under suboptimal conditions or deal with a fluctuating environment. This is a particular problem when scaling engineered yeast cell factories in industrial biotechnology (Takors 2012; Delvigne *et al*. 2014; Wehrs *et al*. 2019). Developing a comprehensive understanding of adaptive responses is a key requirement for the construction of yeast cell factors that are both robust and resilient, and capable of optimal performance in an industrial bioprocess.

Most adaptive responses involve increased or reduced activity of specific proteins, which can be achieved at the level of synthesis, stability or activity. While some specific responses can be at the protein level, for example mediated by allosteric regulation, adaptation usually requires changes in the expression of many genes, and is considered to be a “global” response. Changes in global gene expression can occur at various levels, most notably, via transcription, translation or mRNA stability. Transcriptional changes are due to chromatic restructuring or changes in the activity of particular transcriptional regulators leading to increased or decreased levels of mRNA (Hahn and Young 2011). This is by far the best-understood adaptive process, deployed in response to heat shock (Masser, Ciccarelli and Andréasson 2020), osmotic stress (de Nadal and Posas 2010), oxidative stress (Morano, Grant and Moye-Rowley 2012) and cell wall challenges (Sanz *et al*. 2017; Jiménez-Gutiérrez *et al*. 2020). Transcriptional responses can be studied at the level of individual genes using Northern blots and RT-qPCR, or globally by DNA microarrays or massively parallel sequencing (RNA-Seq), the latter of which has become the method of choice to study changes in gene expression (Schena *et al*. 1995; Gibson, Heid and Williams 1996; Wang, Gerstein and Snyder 2009). Translation results in protein synthesis and, as such, is a better indicator of protein levels than transcription though in many cases, higher abundance of mRNA due to increased transcription leads to a corresponding increase in the amount of translation. This is not always the case, however, and there are also instances where translation is regulated without any changes in the mRNA abundance. A well-documented example of this in yeast is regulation of the translation of the transcriptional activator encoded by *GCN4*. In this case, short open reading frames upstream of the main *GCN4* coding sequence regulate the rate of *GCN4* translation in response to intracellular amino acid levels (Hinnebusch 2005). The TOR growth control system also mediates it some of its effects by regulating translation via controlling access of the small ribosomal subunit to the cap structure at the 5’ end of the mRNA (Merrick 2015). Indeed, as will be mentioned below, there is increased awareness that translational regulation is a central part of the yeast system for controlling gene expression.

Ribosome profiling, sometimes termed Ribo-Seq, is a method that allows for the visualisation and quantification of translation at a global level. First developed in *S. cerevisiae* (Ingolia et al., 2009), it has since been widely used in bacteria, yeast and mammalian systems for genome wide studies of translation (Andreev *et al*. 2017; Ingolia, Hussmann and Weissman 2019; Mohammad, Green and Buskirk 2019). During mRNA translation, a ribosome translocates an mRNA one codon at a time and protects a fragment of mRNA within its mRNA tunnel (Steitz 1969). Ribosome profiling is a method to identify these ribosome protected fragments (RPFs), thereby reporting what mRNAs are being translated at a given point in time. When applying the method, translation is arrested, usually by the addition of translation inhibitors, ribosomes are isolated, and the RPFs identified by deep sequencing. RNA-Seq is usually carried out in parallel to ribosome profiling, allowing estimation of changes in mRNA translation efficiency (Ingolia *et al*. 2009). In *S. cerevisiae*, ribosome profiling has uncovered wide-spread translation of upstream open reading frames (uORFs) (Ingolia *et al*. 2009), non-AUG initiation at canonical genes (Monteuuis *et al*. 2019; Eisenberg *et al*. 2020) and small translated ORFs throughout the genome (Smith *et al*. 2014).

Ribosome profiling combined with RNA-Seq has been useful deciphering both transcriptional and translation regulation in the yeast meiotic programme (Brar and Weissman 2015) and in the response to oxidative stress (Blevins *et al*. 2019). Ribosome profiling has also been carried out on a range of other yeast species including *Saccharomyces paradoxus* (McManus *et al*. 2014), *Schizosaccharomyces pombe* (Duncan and Mata 2014), *Saccharomyces uvarum* (Spealman *et al*. 2018), *Komagataella phaffii* (Alva, Riera and Chartron 2021) and *Candida albicans* (Sharma *et al*. 2021).

We are especially interested in another budding yeast, *Kluyveromyces marxianus*, which originally attracted interest because of its role in food fermentations (Coloretti *et al*. 2017) but is now increasingly being considered as a platform of industrial biotechnology (Fonseca *et al*. 2008; Lane and Morrissey 2010; Karim, Gerliani and Aïder 2020). *K. marxianus* has some intrinsic traits like thermotolerance, a broad substrate range and rapid growth that are useful for biotechnology (Groeneveld, Stouthamer and Westerhoff 2009), and molecular and genomic tools to aids its development as an industrial platform (Cernak *et al*. 2018; Rajkumar *et al*. 2019; Rajkumar and Morrissey 2020). To date, all studies that addressed gene expression in this yeast focused on transcriptional effects via RNA-Seq experiments. Aspects that have been studied include growth and ethanol production on alternative sugar substrates such as xylose (Schabort *et al*. 2016; Kwon *et al*. 2019) and inulin (Gao *et al*. 2015); ethanol tolerance during adaptive laboratory evolution (Mo *et al*. 2019); response to growth inhibitors derived from lignocellulosic substrates (Wang *et al*. 2018) and the ability to grow at high temperatures (Fu *et al*. 2019). Recently, mainly using transcriptome analysis, we determined that young genes specific to *K. marxianus* are enriched in the response to stresses such as high temperature, low pH and high osmolarity (Doughty *et al*. 2020).

To complement the molecular toolbox, and as a resource to study the biology of this yeast, here we report the development of a protocol to carry out ribosome profiling in *K. marxianus.* For this, we adapted and applied the methods previously used for *S. cerevisiae* (Ingolia *et al*. 2009). We also developed a suite of bioinformatics tools to visualise and analyse *K. marxianus* RNA-Seq and ribosome profiling results. This involved addition of the *K. marxianus* data to publicly available genome (GWIPS-viz) and transcriptome (Trips-Viz) browsers, which, in turn, can be uploaded with user-generated expression data and used in private or public configurations. To facilitate the use of ribosome profiling as a very valuable tool to explore gene expression, we also include a detailed step by step protocol for users.

## Materials and Methods

### Strains and growth conditions

*K. marxianus* strain CBS 6556 (CBS-KNAW culture collection, Westerdijk Institute) was used in these studies following standard growth and handling procedures. This particular strain is also available from other collections under the strain name ATCC 26548, NRRL Y-7571, KCTC 17555 and NCYC 2597 and has been quite widely used as a representative *K. marxianus* strain. For ribosome profiling experiments, standard growth conditions used synthetic minimal medium (Verduyn *et al*. 1992) and an incubation temperature of 30°C with shaking. The mineral medium consisted of the following per litre amounts: (NH_4_)_2_SO_4_, 5.0 g; KH_2_PO_4_, 3.0 g; MgSO_4_·7H_2_O, 0.5 g; trace elements (EDTA, 15 mg; ZnSO_4_·7H_2_O, 4.5 mg; MnCl_2_·2H_2_O, 0.84 mg; CoCl_2_·6H_2_O, 0.3 mg; CuSO_4_·5H_2_O, 0.3 ; Na_2_MoO_4_·2H_2_O, 0.4 ; CaCl_2_·2H_2_O, 4.5 mg; FeSO_4_·7H_2_O, 3.0 mg; H_3_BO_3_, 1.0 mg; KI, 0.1 mg); silicone antifoam, 1.5 mL. It was adjusted to pH 6.0 with KOH before autoclaving (121°C, 20 min). The medium was cooled to room temperature and a filter-sterilized solution of vitamins prepared in demineralized water was added, to a final concentration, per liter, of: d-biotin, 0.05 mg; calcium pantothenate, 1.0 mg; nicotinic acid, 1.0 mg; *myo*-inositol, 25 mg; thiamine HCl, 1.0 mg; pyridoxin HCl, 1.0 mg; and para-aminobenzoic acid, 0.20 mg. Glucose was sterilized separately and added to a final concentration of 10 g L^−1^. 150 ml cultures in 500 mL conical flasks were grown to early-log phase at A_600_ ∼0.8 and either harvested or transferred to a shaking water bath at 40°C, with cells harvested at 5, 15, 30 and 60 minutes. All experiments were carried out with two biological replicates.

### Ribosome Profiling

For cell harvesting, cultures were quickly poured into a glass filter assembly (Durapore) with using 0.45 μm pore nitrocellulose filter membrane (GE #7184-009). A vacuum pump was immediately turned on and once liquid media was removed, cells were quickly scraped into a 50 mL falcon tube filled with liquid nitrogen. After harvesting, 1.5 mL of polysome lysis buffer (5 mM MgCl_2_, 150 mM KCl, 20 mM Tris-HCl, 100 µg/mL cycloheximide, 1 mM DTT, 1% Triton X-100) was slowly added dropwise to the liquid nitrogen and cells to create a frozen mixture of buffer and cells. The 50 mL falcon tube (with pierced cap from screwdriver) was placed in -80°C to allow boiling off of the liquid nitrogen. Frozen cells/buffer were disrupted using cryogenic grinding using a Retsch Mixer Mill 400 and 10 mL steel grinding jars and balls. Samples were ground for 6 cycles of 3 minutes each at 20 Hz, the steel jars were submerged in liquid nitrogen to cool samples between each cycle.

After lysis, lysates were gently thawed on ice and quantified with Qubit 4.0 fluorometer and BR-assay kit (#Q10211, Invitrogen). 30 μg of lysate was diluted to 200 μL in polysome buffer (lysis buffer without Triton X-100) and 1.5 μL RNase I was added (Epicentre #N6901K). RNase digestion was carried out at 200 rpm at 37°C for 45 minutes. To halt digestion, SUPERase•In (Invitrogen) was added, and samples were placed on ice before loading onto cold 10-50% sucrose gradients, which were prepared using a Biocomp Gradient Master. Gradients were spun for 3 hours at 4° C and 36,000 RPM (221,632 x g) on SW41-Ti rotor (Beckman Coulter). Monosome fractions were isolated from each sucrose gradient with Brandel Density Gradient Fractionator using 1.5 mL/min flow speed and 60% CsCl, aliquoting fractions every 12 seconds on a UV-visible 96 well plate. Reading the 96 well plate at 260 nm determined which well(s) contained the monosome fractions. RNA from monosome fractions were isolated using Trizol (Invitrogen) (Chomczynski and Sacchi 2006). Ribosome footprints were size selected with a 15% PAGE-Urea gel (70 minutes in 1X TBE and 300 V constant) using a 26 and 34 nt RNA marker (IDT) as a guide for excision. Using a scalpel, a slice representing the RPFs was cut from gel and placed into a 1.5 mL RNase-free Eppendorf tube and 500 μL of RNA elution buffer (300 mM NaOAc pH 5.5, 1 mM EDTA and 0.25% v/v SDS) was added. Following overnight shaking at room temperature to elute the RPFs, RPFs were precipitated using standard alcohol precipitation using ice-cold isopropanol, 80% ethanol and 1.5 μL Glycoblue co-precipitant (Ambion #AM9515).

### Library Construction

In brief, cDNA Library construction with ribosome footprints is based on McGlincy et al. 2017 (McGlincy and Ingolia 2017) protocol with minor modifications. In brief, size selected ribosome footprints were treated with T4 Polynucleotide Kinase (#M0201L, New England Biolabs (NEB)) followed by ligation to a DNA linker using T4 RNA ligase 2, truncated K227Q (#M0351L, NEB). The footprints were reverse transcribed using Protoscript II (#M0368L, NEB). cDNA products were circularized using circligase II (#CL9025K, Epicentre). The major rRNA contaminants were removed using subtractive hybridization with custom biotinylated oligos (Sigma Aldrich) and streptavidin beads (#65001, Invitrogen) as described in Ingolia et al., 2012. The remaining circularized products were amplified by PCR using Phusion polymerase (#M0530L, NEB). In a pilot experiments, libraries were sequenced on MiSeq platform at the Teagasc Next Generation DNA Sequencing Facility, Moorepark, Moorepark West, Fermoy, Co. Cork, Ireland. Prepared libraries using the protocol described within were sequenced on Illumina HiSeq 4000 using SE-75 sequencing at the Genomics & Cell Characterization Core Facility (GC3F), University of Oregon, Eugene, Oregon, USA.

### Ribosome profiling data analysis pre-processing & genome annotation update

Adapter sequences were removed from reads using Cutadapt (Martin 2011). For genomic alignments, rRNA contaminants were removed and remaining reads were aligned to the *K. marxianus* DMKU3-1042 reference genome (Lertwattanasakul *et al*. 2015) with Bowtie (Langmead *et al*. 2009) using parameters -n 2 -m 1. Transcriptome alignments were made with Trips-Viz (Kiniry *et al*. 2019). For all genomic and transcriptome analysis, Ribosome profiling and RNA-Seq reads were aligned to the *K. marxianus* DMKU3-1042 reference strain (Lertwattanasakul *et al*. 2015).

### RNA-Seq

RNA was isolated from clarified lysates using Trizol (Chomczynski and Sacchi 2006) (Invitrogen #15596026) and quantified with a Qubit 4.0 fluorometer (Invitrogen). 1 μg of total RNA from each sample was analysed on an agarose bleach gel to determine RNA quality. Samples were sent to BGI Hong Kong for yeast rRNA removal (Ribo-Zero Gold rRNA Removal kit by Illumina (now discontinued)), library generation and sequencing with paired-end chemistry. Alternatively, RNA-Seq was carried out using polyA selection using Poly(A)Purist Mag Kit (Ambion #AM1922) as per manufacturer’s instructions. PolyA selected cDNA library was generated and sequenced in the same method as ribosome profiling.

## Results and Discussion

### Development of a Ribosome Profiling Protocol to study translation in K. marxianus

The previously-described *S. cerevisiae* protocol (McGlincy and Ingolia 2017) was used as the basis for development of a ribosome profiling procedure for *K. marxianus.* An overview of the pipeline from culturing to downstream bioinformatic analyses is shown in **Fig. 1.** Summarising the first part of the protocol, cultures are rapidly harvested and flash frozen to preserve the translational state of the cell. Frozen cells are then lysed cryogenically in the presence of cycloheximide, which ensures that ribosomes remain stalled even if the cells thaw, and the clarified lysate containing polysomes is treated with RNase I to digest unprotected mRNA surrounding the ribosomes, retaining the ribosome protected fragment (RPF). Monosomes are isolated from a sucrose gradient and loaded on a polyacrylamide gel to allow size selection of RPFs of ∼28nt. A cDNA library of these fragments is created and sequenced to identify the RPFs.

**Figure 1.**
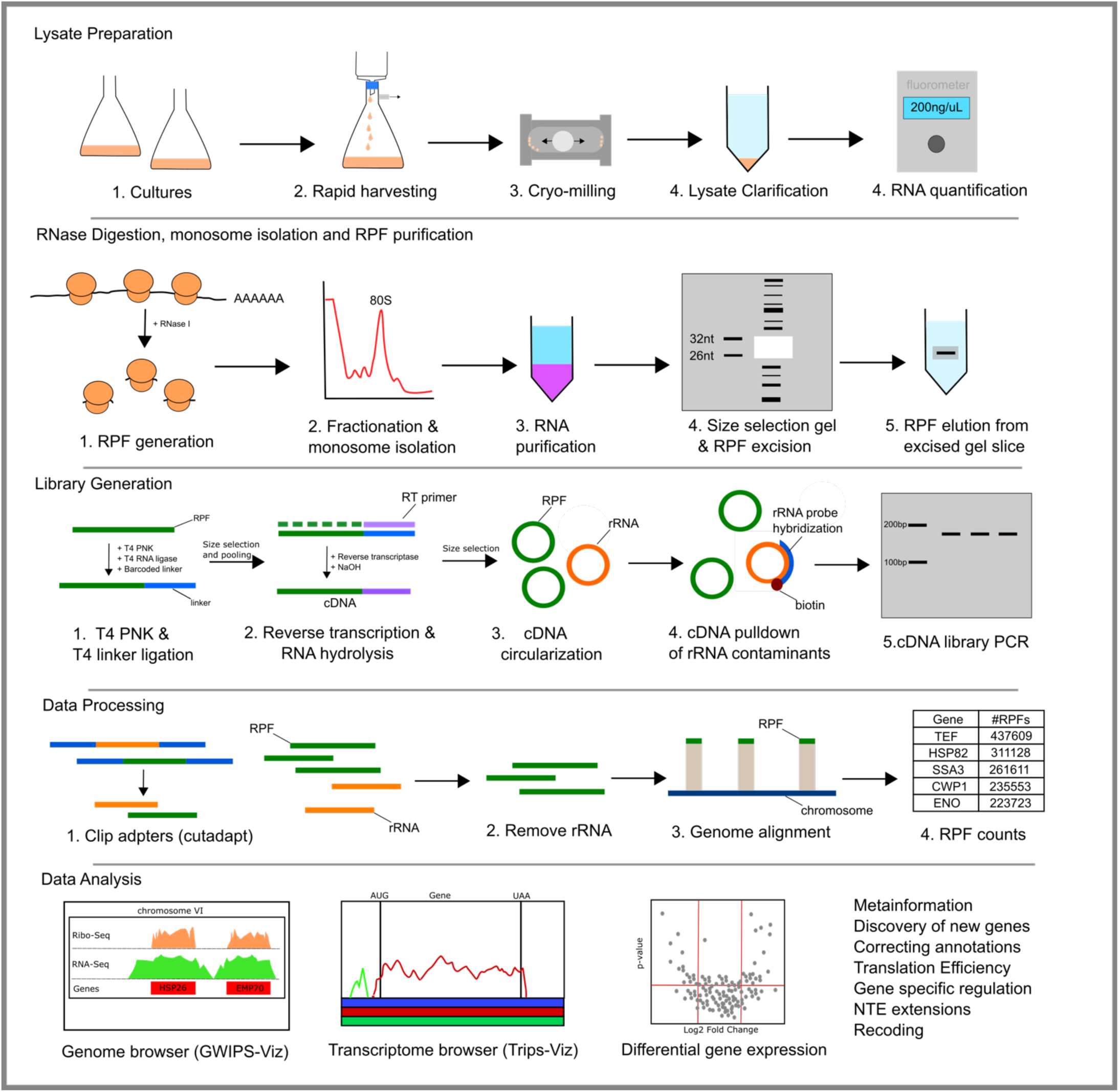
Summary of the ribosome profiling workflow. This summary is broken into five parts including lysate preparation, RPF generation and purification, library generation, data processing and data analysis. Lysate preparation includes culturing, lysis and the quantification of total RNA in a lysate. RNase digestion, monosome isolation and RPF purification represents the generation of ribosome protected fragments (RPFs). Library generation involves the conversion of small RNAs (RPFs) to a cDNA library, ready to be sequenced on an Illumina sequencing platform. Data processing involves removing of the sequencing adapters to leave only RPF sequences which are aligned to the genome. Data analysis typically involves visualizing of data via genome and/or transcriptome browser, differential gene expression and a range of others as listed in figure.

A limited-scale pilot experiment was first carried out to validate the methods and to identify the most abundant rRNA contaminants in the library. These arise because of RNase I digestion of rRNA and subsequent co-purification of fragments of the same size as the RPFs. Due to natural polymorphisms in the rRNA encoding genes between species, the sequence of the major contaminated rRNA fragments needs to be determined empirically for each yeast. Knowing these sequences allows the design of synthetic biotinylated oligos that can be used to reduce rRNA contamination (Ingolia *et al*. 2012). In our pilot library, we identified six highly abundant rRNA fragments, four from the 25S rRNA, and one each from the 18S rRNA and from the mitochondrial 21S rRNA (**Table 1**). One of these sequences from 25S rRNA (GGGTGCATCATCGACCGATCCT) comprised ∼33% of all rRNA contaminants. By reducing rRNA contamination, the proportion of RPFs in a library is increased and thus more usable data are generated per experiment. The ribosome profiling protocol with rRNA depletion was then tested on a larger scale using 150 mL cultures of *K. marxianus* growing at different temperatures to increase the total number of genes that would be expressed. The focus at this time was on assessing the quality of the data generated and the robustness of the protocol rather than on analysis of changes in gene expression. Ribosome profiling was performed in duplicate on flask cultures at 30°C and at 5, 15, 30 and 60 minutes after a transfer from 30°C to 40°C and, as is standard, RNA-Seq was also performed to measure transcript levels. Several analyses were performed to assess the robustness of the data that were obtained. First, the degree of correlation of the number of mapped reads per gene between biological replicates for each condition was assessed and found to be high with a Pearson’s correlation of >0.96 (**Fig. 2A**). Second, we checked whether the RPFs actually represented known protein coding genes (**Fig. 2B**). We found that 14% of reads aligning to the genome represented uniquely mapping RPFs; only ∼0.5% of reads represented ambiguous RPFs, aligning to more than one location on the genome/transcriptome; and ∼85% of the reads mapped to rRNA encoding genes. Third, we examined whether our data showed the distinctive triplet periodicity (or sub-codon phasing) of the aligned reads reflecting the ‘codon-wise’ movement of elongating ribosomes that is seen ribosome profiling data. In the dataset, footprints of length 28nt (approximately half of total footprints) displayed a remarkable strong periodicity signal with ∼95% of RPFs in phase with one of the three sub-codon positions (**Fig. 2C**). Finally, RPFs are expected to be massively enriched in CDS regions of genes. Using metagene profiles, we found that RPFs are largely present with CDS regions (**Fig. 2D and 2E**). In combination, these data demonstrate that the protocol generates robust ribosome profiling data.

**Figure 2.**
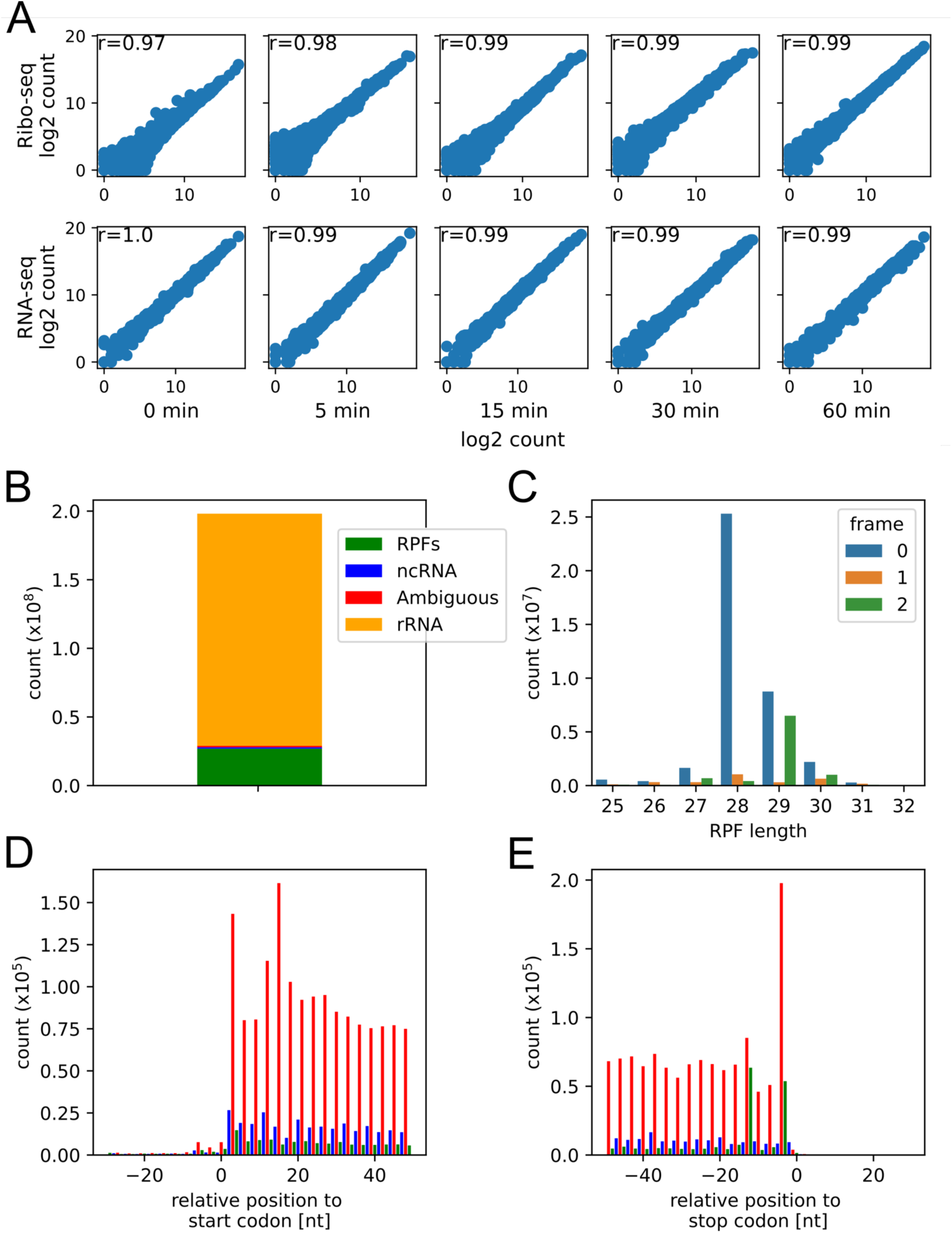
Ribosome Profiling Data from *Kluyveromyces marxianus*. A. Pearson’s correlation of biological replicates for each experimental condition. Axis values represent log2 read counts. B. Composition of ribosome profiling library with rRNA depletion. C. Triplet periodicity of aligned RPFs for each read length. D and E display metagene profile of aligned RPFs near start codon and stop codon, respectively.

**Table 1.**
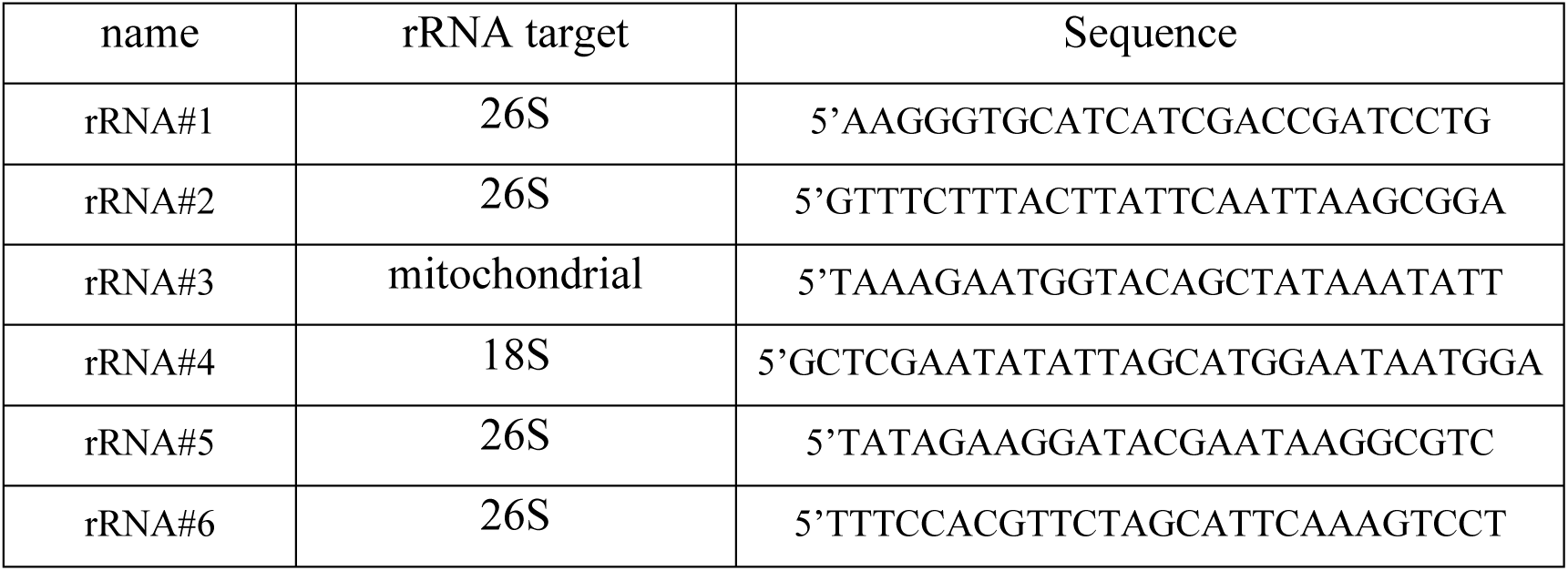
Biotinylated oligos for rRNA depletion. These oligos contain a 5’ biotin modification to allow pulldown of specific rRNA contaminants using magnetic streptavidin beads.

Despite the oligo rRNA depletion, in our dataset from the large-scale experiment, ∼85% of the total reads were rRNA fragments. To determine the efficiency of the targeted rRNA contamination depletion, mapped rRNA contaminant sequences were analysed before and after depletion. After depletion, we see almost 100% efficiency in removal of targeted rRNA contaminants, this is visualised in **Fig.3** where we show efficient reduction of rRNA reads mapping to specific targets of rRNA. It is important to note that the introduction of rRNA contaminants can vary due to slicing of RPFs from size selection gels by free-hand, therefore rRNA abundance and composition may vary between samples and experiments. In our data, we observe this phenomenon where a sequence originating from the 5.8S is present in the post-depletion data but not in the pre-depletion data. If desired, more oligonucleotides could be designed to further reduce rRNA contamination, thus increasing the proportion of RPFs in the sequencing pool. It was interesting to note that while ambiguously mapped reads can represent >10 % of all reads in many studies from *S. cerevisiae* (seen looking at data in the Trips-Viz browser; https://trips.ucc.ie/) these comprised <1% of all reads in *K. marxianus*. Ambiguous mapping ,whereby an RPF maps to two or more loci in the genome or transcriptome, arises because of the very short reads generated by ribosome profiling. As a result, it is not possible to determine the origin of the reads and these are generally discarded/ignored. This difference is most likely due to the large number of paralogous genes in *S. cerevisiae*, which arose through the proposed whole genome duplication/hybridization (WGD) event in the evolutionary history of this species (Wolfe and Shields 1997; Marcet-Houben and Gabaldón 2015). As *K. marxianus* is a pre-WGD yeast, the same issue does not apply.

**Figure 3.**
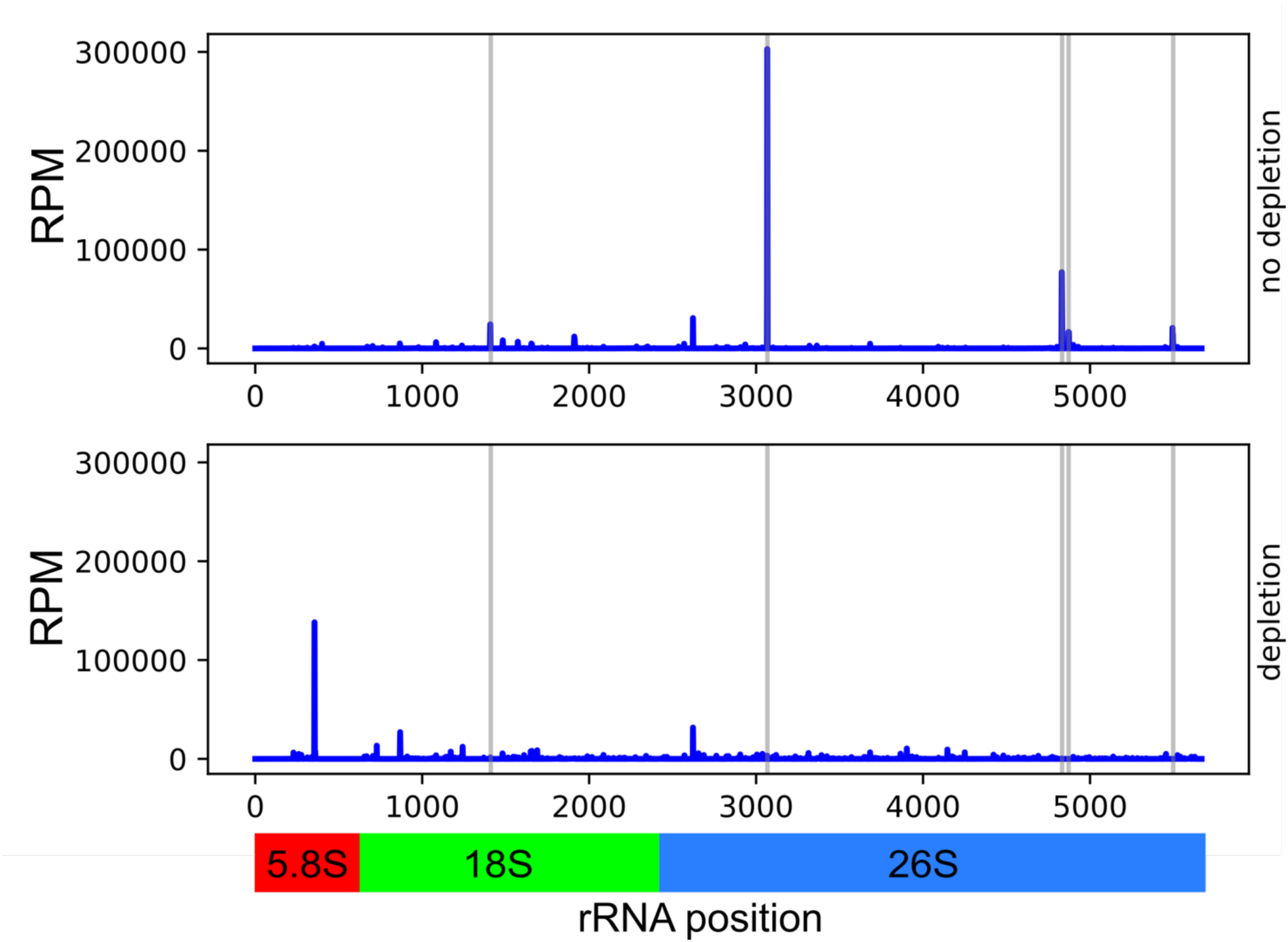
Targeted removal of nuclear encoded rRNA contaminants. Abundance is represented in reads per million (RPM) and position is relative to the generated rRNA index presented in the bottom track. Top panel represents rRNA composition and abundance with no targeted rRNA depletion employed. Bottom panel represents rRNA composition and abundance with targeted rRNA depletion protocol. Targets for rRNA depletion are highlighted as dark grey areas. Abundance is represented in reads per million (RPM) and position is relative to the generated rRNA index presented in the bottom track. The grey vertical lines highlight the rRNA contaminants that are targeted in the oligo depletion step.

### Visualisation of K. marxianus ribosome profiling data on public browsers

Visualisation of ribosome profiling data is important to examine translation/transcription of particular loci of interest. We previously developed two tools to allow visualisation of these data at a genome level (GWIPS-viz) and at the level of individual RNAs (Trips-Viz). These tools are freely accessible via RiboSeqOrg portal at https://riboseq.org. GWIPS-viz is a genome browser that displays RPFs mapped to each chromosome of a reference genome (Michel *et al*. 2014). The GWIPS-viz database already contained reference genomes for ∼24 animal, plant, protozoal, fungal and viral genomes and we added *K. marxianus* using the genome sequence and annotation from *K. marxianus* DMKU3-1042 strain as this was the most complete genome sequence available (Lertwattanasakul *et al*. 2015). It is possible to search GWIPS-viz by gene name or gene ID and to zoom in / out of loci and as an extra feature that is new to GWIPS-viz, we included strand orientation of our ribosome profiling data to allow users determine the strand to which a RPF is mapped (orange for +/forward strand, blue for -/negative strand) (**Fig. 4**). The browser is free to use and any user that generates their own ribosome profiling data or RNA-Seq tracks (bigWigs) can upload those data as custom tracks that can be viewed privately or made public. Once uploaded, a user is able to visualise and analyse their data using all the functionality of GWIPS-viz.

**Figure 4.**
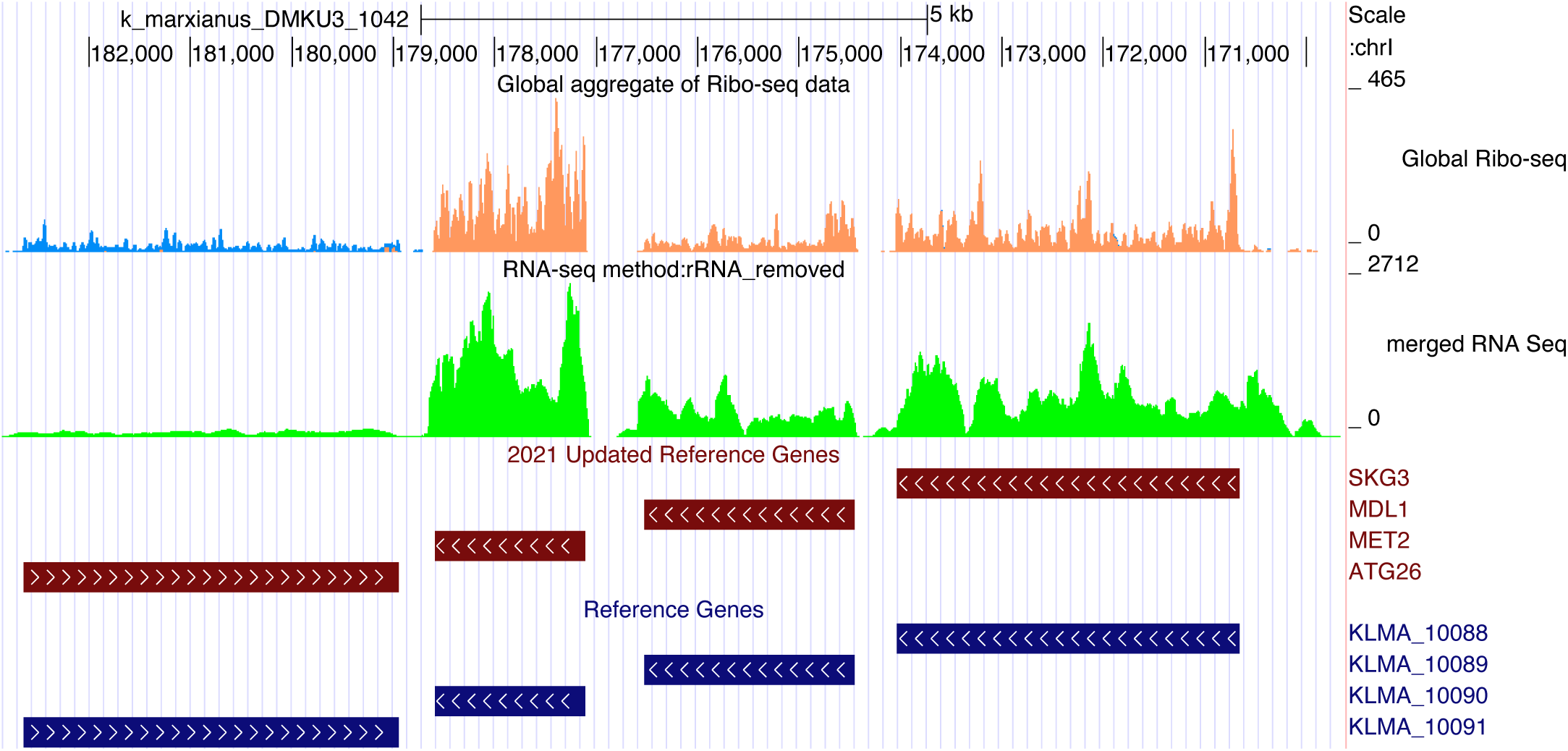
GWIPs-Viz Browser screenshot surrounding the *SKG3*, *MDL1*, *MET2* and *ATG26* locus of chromosome 1. Arrows on Reference Gene bars represent strand orientation. For Ribosome Profiling, orange reads represent positive strand RPFs while blue reads represent negative strand RPFs.

GWIPS-viz is mainly designed for analysis at a global level, allowing users to visualise any part of a genome, regardless of whether or not it is included in the annotations. In contrast, a second tool Trips-Viz, is a transcriptome level browser that focuses on individual mRNAs and allows a deep analysis of translation of each mRNA (Kiniry *et al*. 2019, 2021). This transcriptome browser allows users to generate single transcript plots displaying the open reading frame that is being translated. It also allows users to visualise the distribution of RPFs along an individual mRNA while also utilising the triplet periodicity signal and differential colouring to identify potential translation in each open reading frame. As Trips-Viz did not include a reference transcriptome for *K. marxianus*, we created this reference transcriptome using our data. The application of Trips-Viz to study an individual mRNA is illustrated with an analysis of *HSP26*, using the (ribosome profiling) data for translation at 30°C and 40°C (**Fig. 5**). The top panel uses aggregate data and shows the distribution of RPFs between each reading frame and across the transcript. It is clear that reads from the first open reading frame (red) dominate, which match the position and frame of the annotated CDS. The increase in the number of reads (RPFs) at certain positions indicates ribosome stalling during translation; for example, at difficult to translate codons. The bottom panel compares the normalised read count of the correct open reading frame between the samples coming from cells grown at 30°C and 40°C. The huge increase in translation at 40°C is evident. This was to be expected as *HSP26* encodes a heat shock protein and is strongly transcriptionally induced by temperature shift. Thus, the increase in translation in this case is due to an increase in mRNA abundance. Although not shown in this simple example, in addition to single transcript plots, the Trips-Viz browser contains a large amount of metadata analyses such as triplet periodicity, read breakdowns, metaplot, protein count tables, differential expression analyses that is useful for detailed studies of translation and its regulation.

**Figure 5.**
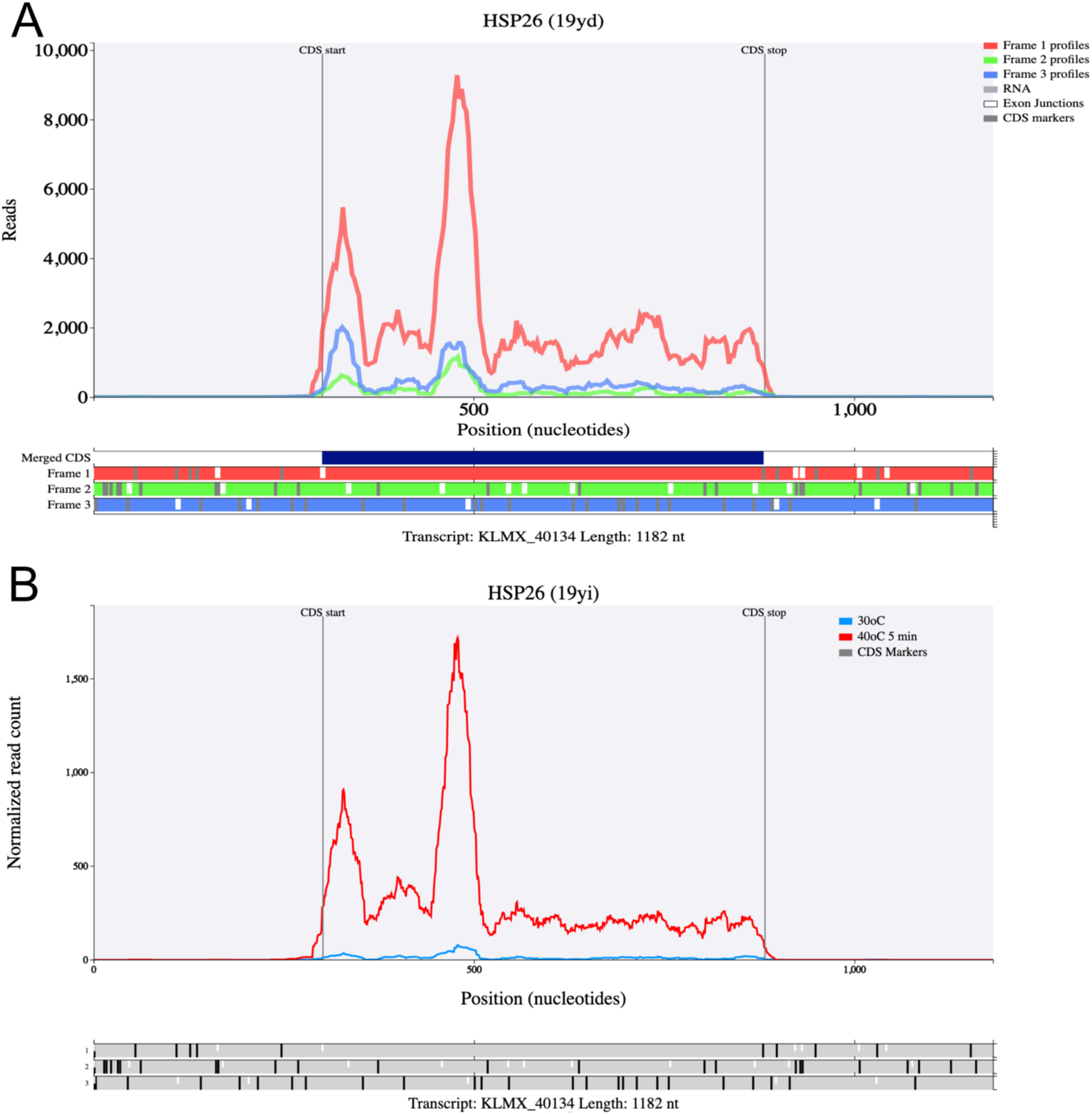
Trips-Viz transcript plots of *HSP26*. A. Ribosome profiling coverage for aggregated data is displayed. Note the dominant red line corresponds to the *HSP26* CDS region. B. Normalized transcript comparison plot of HSP26 showing increased mRNA translation at 30°C and 40°C temperatures.

### Integrated omics studies with K. marxianus

Ultimately, a full suite of omics technologies, ranging for genomics to proteomics, is a requisite for comprehensive studies of any microbe. Analysis of genome sequences and transcriptomes is now relatively straightforward for diverse yeasts, but the development of other tools still lags. Now, with the laboratory and *in silico* methods that we developed, it is possible to perform ribosome profiling with a non-model yeast, *K. marxianus*. By including both transcriptome (RNA-Seq) and translatome (ribosome profiling) analysis in future studies, it will be possible to generate a comprehensive view of gene expression at a point of time and in response to a perturbation. This can be very useful to understand biological processes and for the development of strains for biotechnology. The strategy taken, and the pipeline used, can also serve as a prototype for the development of ribosome profiling methods for other yeast of biological and biotechnological interest. To facilitate the application of the tools by as many users as possible, a comprehensive step by step protocol is provided as a supplementary protocol.

## Data Accessibility

Ribosome profiling and RNA-Seq datasets have been deposited to the European Nucleotide Archive under the under the project accession number PRJEB45612. The data have also been deposited to GWIPS-viz https://gwips.ucc.ie/ and to Trips-Viz https://trips.ucc.ie/.

## Acknowledgements

We greatly appreciate the help of Dr Gary Loughran for advice on developing ribosome profiling and Luke Power (RiboMaps) for his critical reading of the manuscript. We thank Dr Simon Lawrence, School of Chemistry, UCC, for providing access to the Retsch mixer mill and Matthew Leibovitch (McGill University) for his help installing the tracerDaq hardware. This project received funding from the European Union’s Horizon 2020 Framework Programme for Research and Innovation Grant Agreement No. 720824. This work was also supported by SFI-HRB-Wellcome Trust Biomedical Research Partnership [210692/Z/18/ to P.V.B.].

## Materials

### Strains

*Kluyveromyces marxianus* CBS 6556 (ATCC 26548/ NCYC 2597/ KCTC 17555).

### Equipment

- Thermocycler (Applied biosystems 2720 Thermal Cycler)
- Nutator (Stuart see-saw rocker SSL4)
- Shaker (Heidolph Titramax 1010)
- Vortex Genie 2 (Scientific Industries)
- Refrigerated centrifuge
- Beckman ultracentrifuge and SW41ti swinging bucket rotor
- Ultracentrifuge polypropylene tubes (14 x 89 mm) (Beckman #331372)
- Ultracentrifuge polypropylene tubes (8 x 34 mm) (Beckman #343777)
- T-shaker (thermo-shaker for 1.5 mL tubes, T-shaker EuroClone)
- Retsch Mixer Mill (model: mm200)
- Retsch 10 mL stainless steel grinding jars and grinding balls
- Heat block (Stuart block heater SBH130D)
- Qubit Fluorometer 4.0
- Qubit assay tubes
- Micropipettes and filter tips (P2, P10, P20, P200 and P1000).
- RNase-free PCR tubes
- 1.5 mL RNase-free tubes
- 2.0 mL RNase-free tubes
- 15 mL RNase-free falcon tubes
- 50 mL RNase-free falcon tubes
- Gradient master (BIOCOMP #B108-2)
- Density Gradient Fractionation System (Brandel BR-188)
- TracerDaq USB-1208FS (Measurement Computing)
- UV Plate Reader (Spectramax M3, Molecular Devices)
- 0.45 μM Whatman cellulose nitrate filter membranes (GE life sciences #7184-004)
- Steriflip Sterile 50 mL Disposable Vacuum Filtration System 0.22μm (Millipore #SCGP00525)

### Enzymes

- RNase I 10 U/µL (Lucigen #N6901K)
- T4 PNK 10 U/µL (NEB #M0201L)
- T4 RNA Ligase 2, truncated K227Q 200 U/µL (NEB #M0351L)
- Protoscript II 200U/µL (NEB #M0368L)
- 5’ DNA Adenylation Kit (NEB #E2610L)
- CircLigase II 100U/µL (Epicentre #CL9025K)
- Phusion Polymerase 2U/µL (NEB #M0530)

### Reagents

- Ultra-pure water
- Yeast Extract (Sigma-Aldrich #Y1625)
- Tryptone (Fluka analytical #9410)
- D-(+)-Glucose (Sigma-Aldrich #G7021)
- Cycloheximide ≥95% (HPLC) (Sigma-Aldrich #01810)
- SUPERase•In™ RNase inhibitor (Invitrogen #AM2696)
- PEG-8000 (supplied with T4 RNA ligase 2, truncated K227Q).
- dNTPs 10mM (NEB #N0447L)
- Streptavidin C1 Dynabeads (Invitrogen #65001)
- ssRNA low range ladder (NEB #N0364S)
- Oligo Clean & Concentrator kit (Zymoresearch)
- Qubit BR-RNA assay kit (Life Technologies #Q10210)
- Glycoblue coprecipitant (Ambion #AM9515)
- Oligo clean and concentrator (Zymoresearch #D4060)

### Chemicals

- Magnesium Chloride (Sigma-Aldrich #M2670)
- Sodium chloride (Sigma-Aldrich #S7653)
- Sodium Hydroxide (Sigma-Aldrich #S8045)
- Trizol (Invitrogen #10296010)
- Sucrose (Sigma-Aldrich #S0389-500g)
- Ethanol (Sigma-Aldrich #51976)
- Chloroform (Sigma-Aldrich #C2432)
- Isopropanol (2-propanol, Sigma-Aldrich #I9516)
- Triton X-100 (Sigma-Aldrich #X100)
- DTT (Sigma-Aldrich #D9779)
- Potassium chloride (Sigma-Aldrich #P9541)
- Tris Base (Fisher Scientific #CAS 77-86-1)
- Urea (Fisher Scientific #BP169-500)
- APS (Sigma-Aldrich #A3678)
- Cesium Chloride (Sigma-Aldrich #C4036-1kg)
- TEMED (Sigma-Aldrich #T9281)
- EDTA (Sigma-Aldrich #EDS)
- Sodium dodecyl sulfate, SDS (Sigma-Aldrich #L4509)
- NaOAc (Sigma-Aldrich #S8625)
- 40% acrylamide/bis-acrylamide (19:1) (Sigma-Aldrich #A9926)
- Liquid nitrogen

## Oligonucleotides

Oligonucleotides used in this study (same as McGlincy and Ingolia, 2017) were purchased from Integrated DNA Technologies (IDT) except for the biotinylated oligos for rRNA depletion which are sourced from Sigma.

### RPF markers

- These are ordered as RNA oligonucleotides.
- The 3’ ends of these oligos are phosphorylated, to allow these to be used as controls for dephosphorylation (with T4 PNK) and subsequent linker ligation.

32mer marker NI-800: 5′-AUGUACACUAGGGAUAACAGGGUAAUCAACGCGA/3Phos/

26mer marker, NI-801: 5′-AUGUUAGGGAUAACAGGGUAAUGCGA/3Phos/

### Linkers

- These are ordered as DNA oligonucleotides.
- Each linker contains a unique 5 nt barcode to allow pooling of ligations before cDNA library preparation.

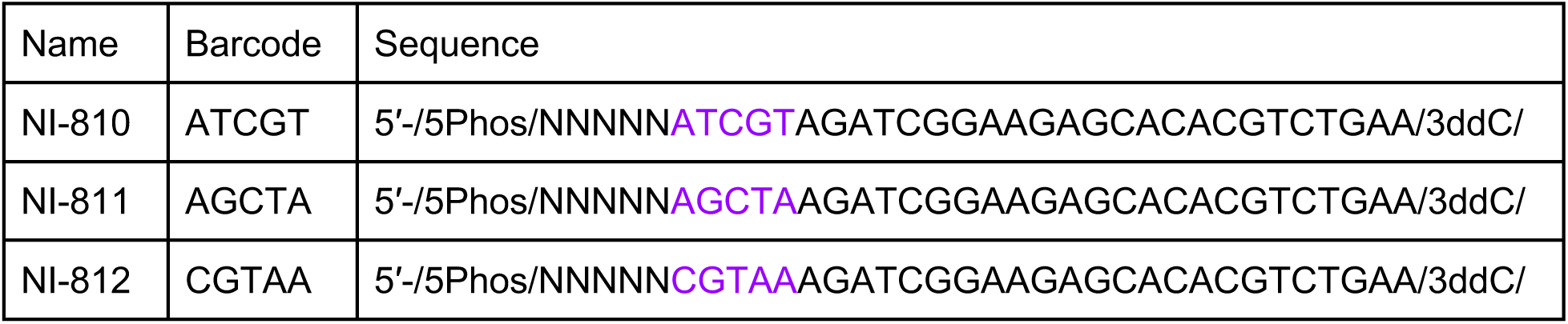
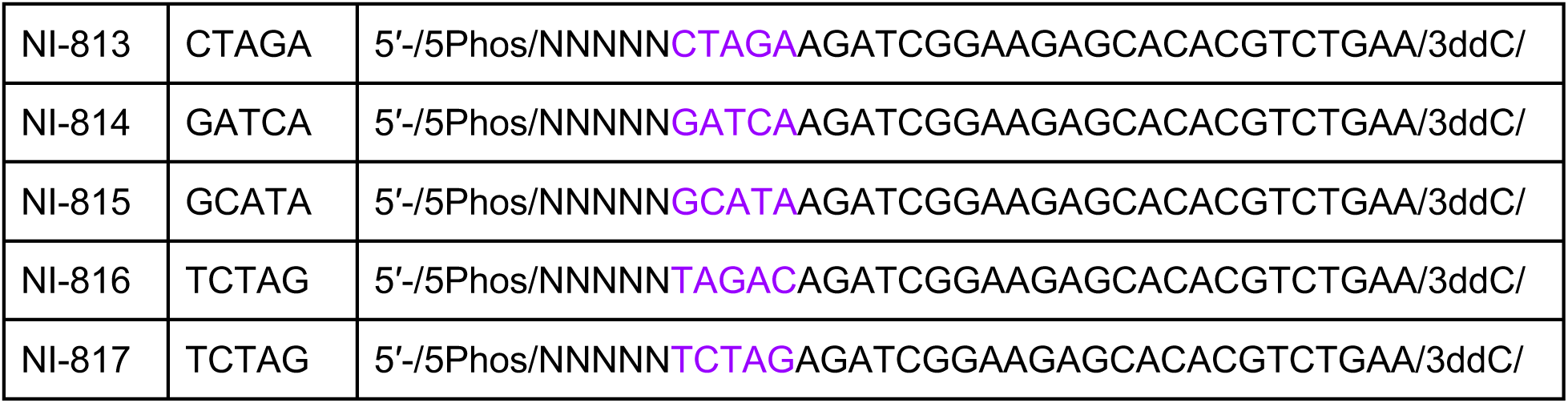

### RT primer

- The RT primer is ssDNA oligonucleotide, containing an 18-atom spacer modification (iSp18). This spacer is required during the final library PCR to amplify the library from circularized cDNA.
- Also included are 2 random nucleotides at the 5’end of RT primer.

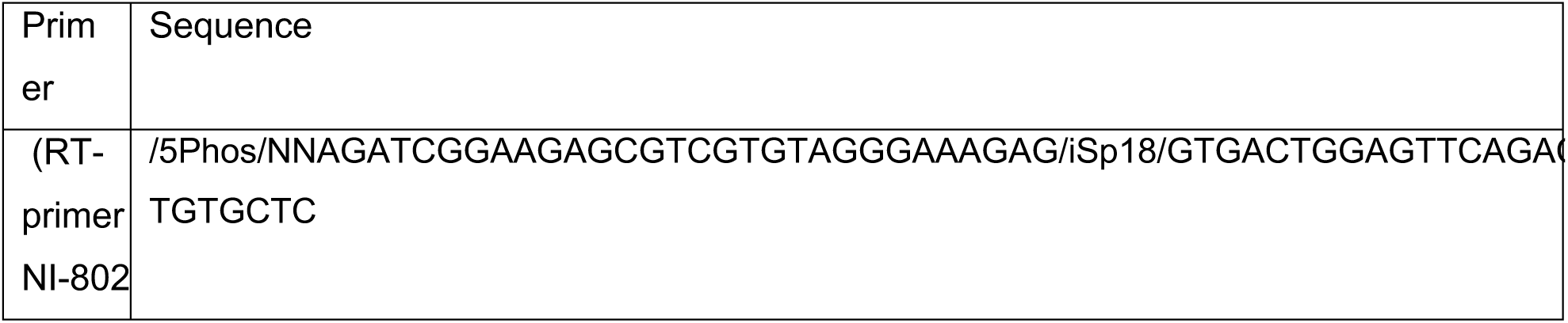

### Biotinylated oligos for rRNA depletion

- These DNA oligonucleotides are biotinylated at the 5’ends and are purified via HPLC.

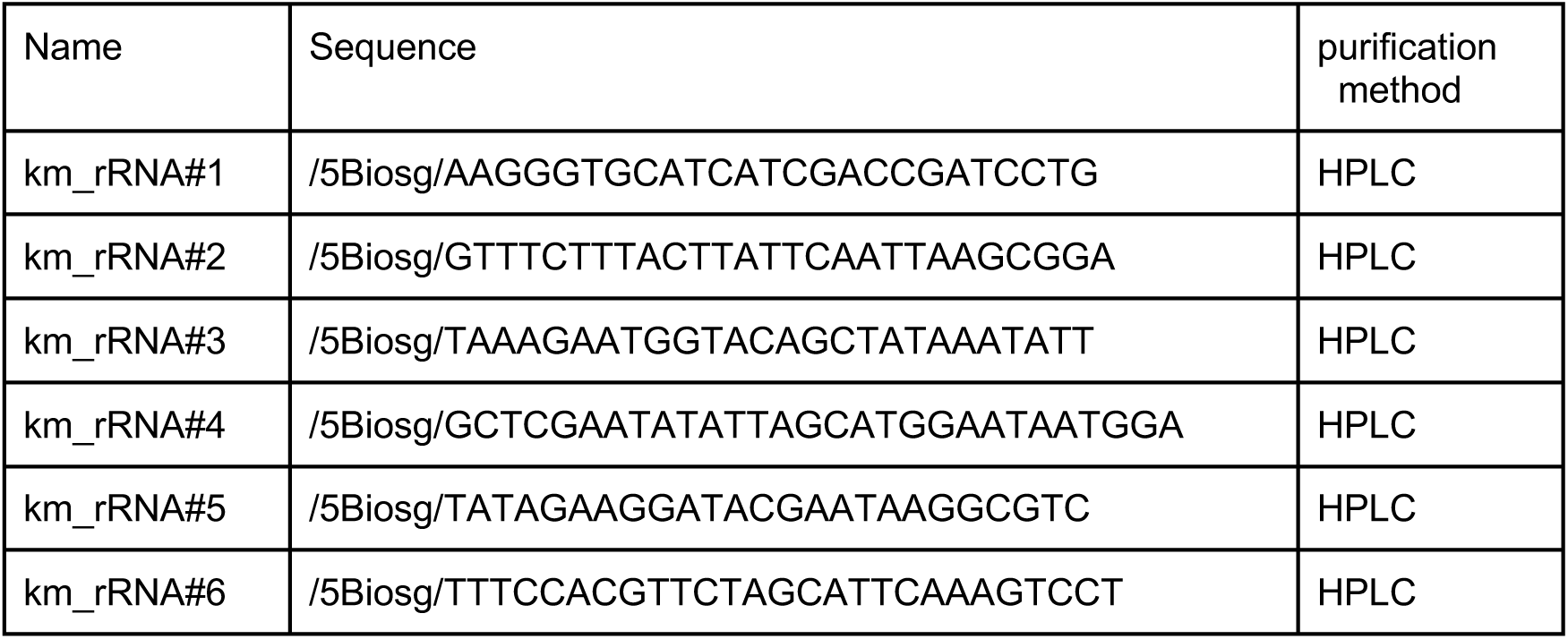

### cDNA library PCR primers

- The Illumina forward primer is the same in each reaction while each sample includes different reverse primer to allow multiplexing.
- Each reverse primer includes a unique barcode which allows for multiplexing of samples as follows, whereby JJJJJJ indicates a unique barcode:

5′- CAAGCAGAAGACGGCATACGAGATJJJJJJGTGACTGGAGTTCAGACGTGTG

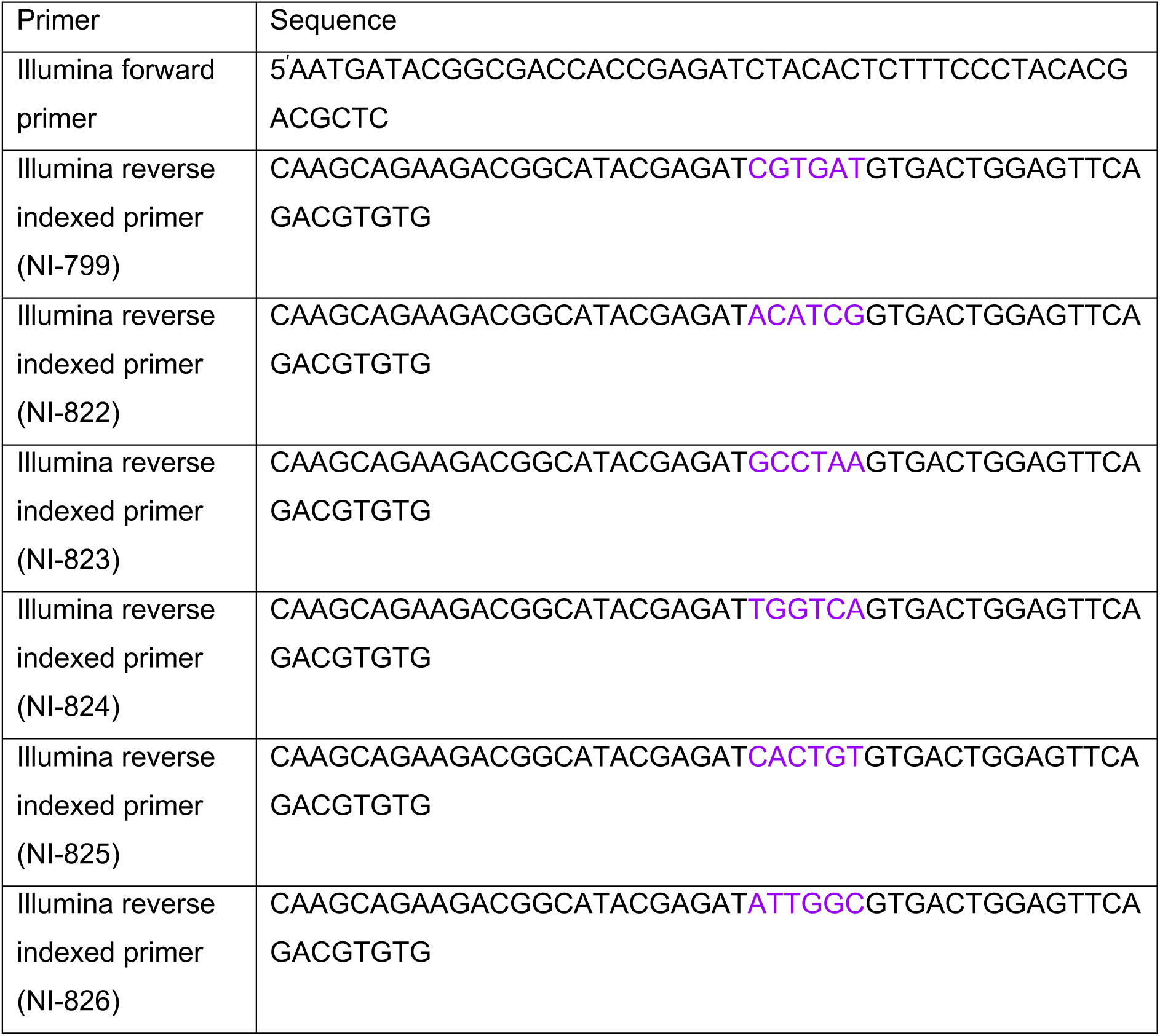

## Solutions/Buffers

### Polysome lysis buffer and Polysome buffer

- Keep on ice.
- Prepare this buffer excluding Cycloheximide (CHX) and DTT, freeze at -20°C. On day of use, thaw on ice and add Cycloheximide from a 100 mg/mL in DMSO stock and DTT, keep on ice.
- For polysome buffer, substitute Triton X-100 with water as Triton X-100 is only required for lysis/lysate preparation.

| Component | Volumes | Final Concentration |
| --- | --- | --- |
| 1M Tris-HCL 7.5 | 200 µL | 20 mM |
| 2M KCl | 750 µL | 150 mM |
| 1M MgCl<sub>2</sub> | 100 µL | 10 mM |
| Cycloheximide (100mg/ mL) | 10 µL | 100 µg/ mL |
| 1M DTT | 10 µL | 1 mM |
| 20% Triton X-100 | 500 µL | 1% |
| Ultra-pure water | 8.43 mL |  |
| Total Volume | 10 mL |  |

### Polysome gradient solutions

- Make polysome gradients fresh.
- First make a polysome gradient buffer (without sucrose) which is added to a 50 mL falcon tube containing the required weight of sucrose.
- Once buffer is added to the sucrose, vortex each solution rigorously until sucrose is dissolved.
- The 10% and 50% sucrose solutions can be left on a nutator for 10 minutes to reduce bubbles in the tubes and leave on ice until use.

1. Polysome Gradient Buffer

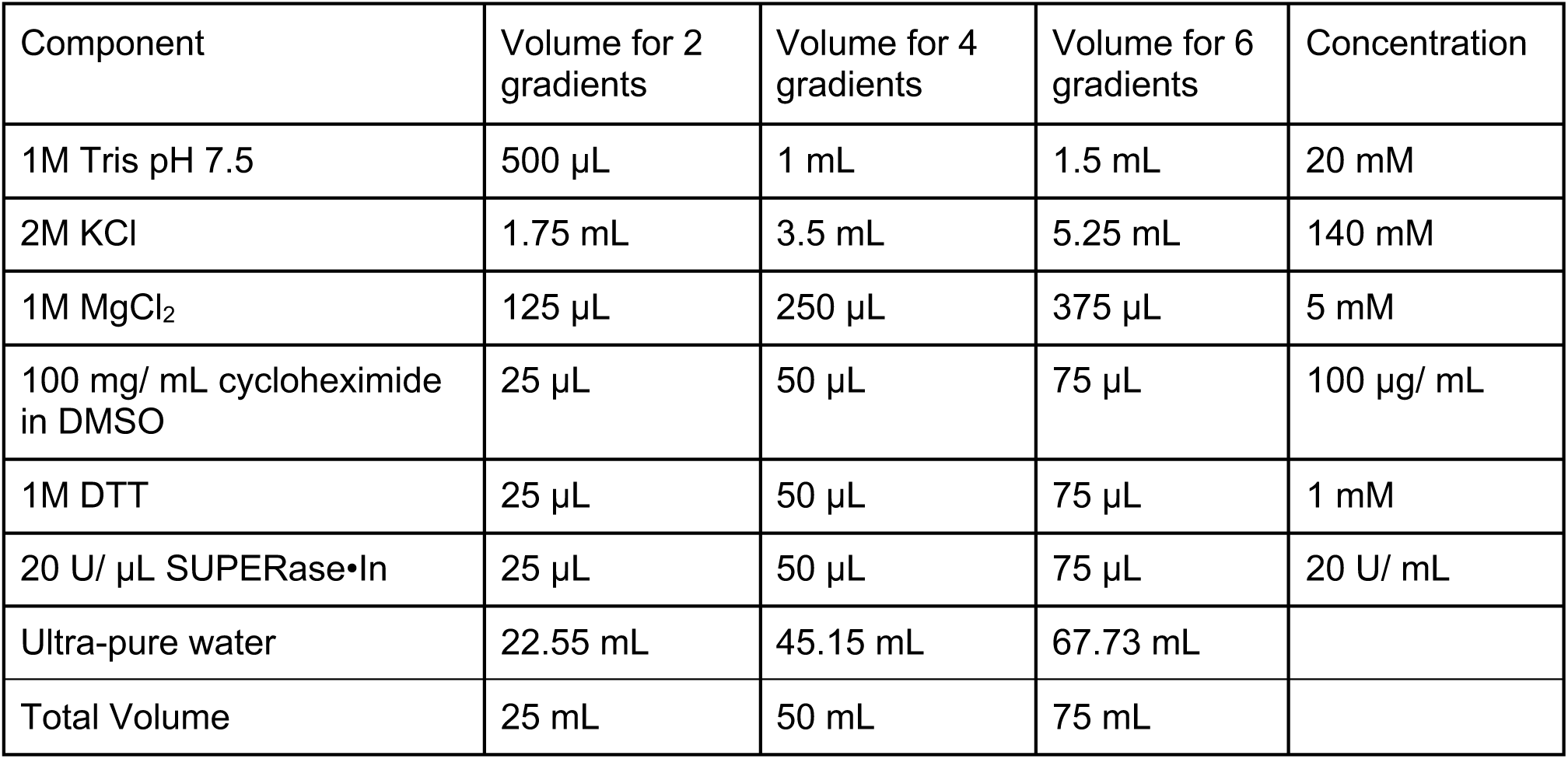

| Component | Volume for 2 gradients | Volume for 4 gradients | Volume for 6 gradients | Concentration |
| --- | --- | --- | --- | --- |
| 1M Tris pH 7.5 | 500 μL | 1 mL | 1.5 mL | 20 mM |
| 2M KCl | 1.75 mL | 3.5 mL | 5.25 mL | 140 mM |
| 1M MgCl<sub>2</sub> | 125 μL | 250 μL | 375 μL | 5 mM |
| 100 mg/ mL cycloheximide in DMSO | 25 μL | 50 μL | 75 μL | 100 μg/ mL |
| 1M DTT | 25 μL | 50 μL | 75 μL | 1 mM |
| 20 U/ μL SUPERase·In | 25 μL | 50 μL | 75 μL | 20 U/ mL |
| Ultra-pure water | 22.55 mL | 45.15 mL | 67.73 mL |  |
| Total Volume | 25 mL | 50 mL | 75 mL |  |

2. 10% Sucrose (w/v):

| Component | Amount for 2 gradients | Amount for 4 gradients | Amount for 6 gradients |
| --- | --- | --- | --- |
| RNase-free sucrose | 1.5 g | 3 g | 4.5 g |
| Polysome gradient buffer | 13.9 mL | 27.8 mL | 41.7 mL |

3. 50% Sucrose (w/v):

| Component | Amount for 2 gradients | Amount for 4 gradients | Amount for 6 gradients |
| --- | --- | --- | --- |
| RNase-free sucrose | 7.45 g | 14.9 g | 22.35 g |
| Polysome gradient buffer | 10.4 mL | 20.8 mL | 31.2 mL |

### 60% CsCl Solution

To make 100 mL of 60% (w/v) CsCl:

1. Weigh 60g of CsCl.
2. Add ultra-pure water to 100 mL.
3. Vortex/shake until CsCl solution is fully dissolved.
4. Stain the solution with bromophenol blue by dipping the tip of a metal spatula into bromophenol blue powder and into the bottle of 60% CsCl until the solution becomes clear blue. Solution can be stored at room temperature.

### Recipe for 15% PAGE gel with Bis-Acrylamide and Urea

- Note that 15% Urea-PAGE gel solution must be prepared without 10% APS and TEMED, these two chemicals are added immediately before making the gel.
- Stocks of this Urea-PAGE gel solution (without 10% APS and TEMED) can be prepared and stored at RT for 3 months. Protect PAGE solutions from light, store in dark cupboard or cover with foil.
- For all PAGE-gels, add 10% APS and TEMED to stock solutions in a 15 mL falcon tube, vortex briefly and immediately pour into a 0.75 mm gel cassette with a 10 mL pipette.

For a stock solution, add the following to an appropriately sized tube/bottle. Dissolve the solution in a 37°C water bath (vortexing during the incubation can speed up the dissolving process). After the solution is dissolved, filter with steriflip and store at RT.

| Component | Stock for 1 gel | Stock for 2 gels | Stock for 6 gels |
| --- | --- | --- | --- |
| 40% acrylamide/bis acrylamide (19:1) | 2.815 mL | 5.63 mL | 18.76 mL |
| Urea | 3.6 g | 7.2 g | 24 g |
| 10X TBE | 750 µL | 1.5 mL | 5 mL |
| Ultra-pure water | 950 µL | 1.9 mL | 6.33 mL |
| Total volume | 7.5 mL | 15 mL | 50 mL |

### 3X RNA loading dye

3X RNA loading dye is composed of 6M Urea, 25% sucrose and 1-2 grains of bromophenol blue.

### RNA extraction buffer

RNA extraction buffer is composed of 300 mM NaOAc, 1mM EDTA and 0.25% SDS (v/v). Store at room temperature.

| Component | Volume | Final Concentration |
| --- | --- | --- |
| 3M NaOAc pH 5.5 | 1 mL | 300 mM |
| 0.5M EDTA | 20 μL | 1 mM |
| 10% SDS | 250 μL | 0.25% v/v |
| Ultra-pure water | 8.73 mL |  |
| Total volume | 10 mL |  |

### Recipe for 7.5% Urea-PAGE gel with Bis-Acrylamide and Urea

- This 7.5% Urea-PAGE gel is used for cDNA size selection after reverse transcription.
- Volumes for larger stock solutions are shown below (15 mL or 50 mL) which can be stored at RT (protected from light), adding 10% APS and TEMED immediately before gel loading.

Prepare the following Urea-PAGE gel solution.

|  |  |  |  |
| --- | --- | --- | --- |
| 40% acrylamide/bis-acrylamide (19:1) | 1.41 mL | 2.82 mL | 9.4 mL |
| Urea | 3.6g | 7.2g | 24g |
| 10X TBE | 0.75 mL | 1.5 mL | 5 mL |
| Ultra-pure water | 2.36 mL | 4.72 mL | 15.73 mL |
| Total volume | **7.5 mL** | **15 mL** | **50 mL** |

Dissolve in a water bath at 37°C then filter with steriflip. Store at RT.

7.5 mL Gel = 7.48 mL stock + 18.75 μL 10% APS + 3.75 μL TEMED

### DNA Extraction Buffer

DNA extraction buffer is composed of 300 mM NaCl, 1 mM EDTA and 10 mM Tris pH 8.0. Store at room temperature.

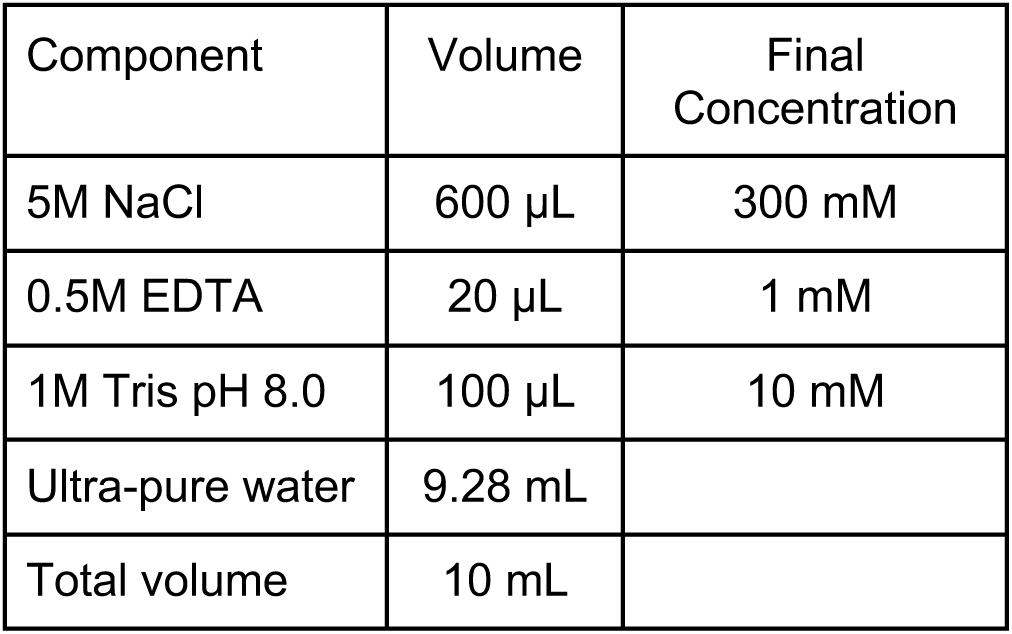

| Component | Volume | Final Concentration |
| --- | --- | --- |
| 5M NaCl | 600 μL | 300 mM |
| 0.5M EDTA | 20 μL | 1 mM |
| 1M Tris pH 8.0 | 100 μL | 10 mM |
| Ultra-pure water | 9.28 mL |  |
| Total volume | 10 mL |  |

### 20X SSC (hybridization buffer)

- A 20X stock solution contains 3 M sodium chloride and 300 mM sodium citrate adjusted to pH 7.2, may be stored at room temperature.

For 100 mL stock solution (scale down if required):

1. Weigh out 17.53 g NaCl and 8.82 g Sodium Citrate.
2. Add 80 mL ultra-pure water and dissolve.
3. Adjust pH to 7.2 with HCl.
4. Bring the final volume to 100 mL.

### 8% polyacrylamide gel for cDNA library products

- This 8% PAGE gel does not require urea and is used for dsDNA products from the final library PCR stages.
- Add 10% APS and TEMED last, allow 30 minutes to polymerize at room temperature.

| Component | 8% |
| --- | --- |
| 10X TBE solution | 750 μL |
| 19:1 acrylamide:bisacrylamide solution (40%) | 1.5 mL |
| Water | 5.25 mL |
| TEMED | 15 μL |
| Ammonium Persulfate (10% w/v) (APS) | 52.5 μL |
| Total Volume | 7.57 mL |

**Figure S1.**
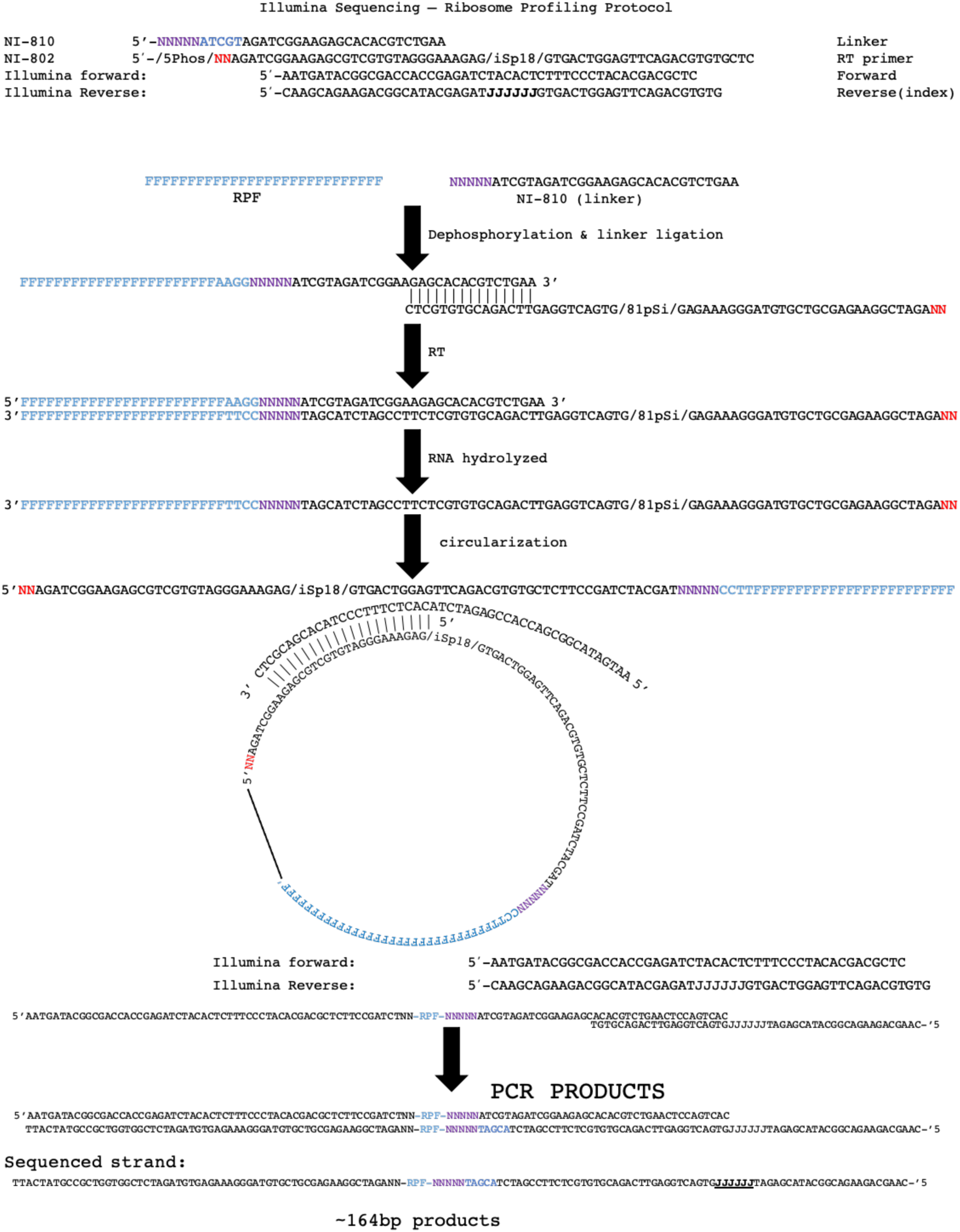
Molecular schematic of ribosome profiling library preparation. Oligos used in library generation are displayed at top of figure.

## Protocol for ribosome profiling

### Safety Precautions

While handling liquid nitrogen and chilled material, wear appropriate safety glasses and PPE, use cryogenic gloves to avoid burns, and adhere to all lab safety procedures.

#### 1.1 Growth

- *Kluyveromyces marxianus* strain CBS 6556 (CBS-KNAW culture collection, Westerdijk Institute) was routinely used in this protocol but the method should work equally well with other strains. In the ribosome profiling data shown in this study, cultures were grown to early-log phase at A_600_ 0.6 - 0.8 in 150 mL synthetic minimal medium (Verduyn *et al*., 1992) in 500 mL conical flasks with shaking.

1. 5 mL of YPD or synthetic minimal medium (Verduyn *et al*., 1992) in a sterile 50 mL falcon tube is inoculated with a fresh colony of yeast and incubated overnight at 30°C in a shaking incubator. Next day, transfer an appropriate volume of overnight culture to 150 mL media in a 500 mL Erlenmeyer flask so that the A_600_ is diluted to ∼0.06.
2. Incubate the culture in a shaking incubator at 30°C (or other) to ∼0.6 – 0.8 A_600_ (log/exponential-phase). As the cells reach this desired OD600, prepare harvesting apparatus in following section.

#### 1.2 Harvesting

- It is important to implement rapid harvesting of cells for ribosome profiling experiments. If cells are left for an extended period on exposed filter membrane, it may induce stress which can quickly alter global translation and compromise the experiment.
- In this protocol, a glass filter apparatus is used to rapidly remove media and trap cells on a filter membrane. The cells are immediately scraped off the membrane, flash frozen and stored in liquid nitrogen.
- The rapid filtration takes us ∼10-12 seconds from pouring the culture onto the membrane and placing the scraper into the liquid nitrogen.

1. Before harvesting of cells, prepare the following equipment:

a. Assemble the glass filter assembly and filter membrane.
b. Prepare a 50 mL falcon tube for each sample by piercing the cap with a screwdriver to make a single hole which allows the evaporation of liquid nitrogen. **IMPORTANT:** make sure to pierce the cap, failing to do so will result in tube explosion.
c. Place the falcon tube into a rack inside a styrofoam box and fill the falcon tube 3/4 with liquid nitrogen.
2. Quickly pour the 150 mL culture into the glass filter assembly and turn on the vacuum.
3. As the media becomes drained, have a cell scraper ready and when media is drained, quickly remove the clamp/upper glass funnel and scrape the cells off the membrane, scrape the membrane enough so that all of the membrane becomes scraped at least once. A clear visible accumulation of yeast on the scraper will be present.
4. Immediately place the cell scraper into the liquid nitrogen-filled falcon tube and leave it for 30 seconds to freeze. Using another pre-chilled cell scraper, detach the cell pellet from the original scraper. If processing multiple samples in quick succession proceed to step 6 and when all samples are collected, remove tubes from -80°C, add liquid nitrogen to the pellet and proceed to step 5.
5. Using a P1000, slowly add 2 mL of polysome lysis buffer (PLB) to the falcon tube so that the PLB forms frozen droplets alongside the cell pellet, add more liquid nitrogen if needed.
6. Place the pierced cap on the tube and place upright in -80°C for storage. The excess liquid nitrogen will boil off through the pierced cap. The frozen cells can be stored for extended periods.

#### 1.3 Lysis

- Here, cryogenic mechanical lysis is used to break the cells and release ribosomes/polysomes.
- Cryo-milling ensures ribosomes are unable to translocate and reduces RNA degradation and upon thawing ribosomes are exposed the cycloheximide which will reduce translocation.

1. Place the unscrewed grinding jars and grinding balls into a liquid nitrogen filled styrofoam box. Add enough liquid nitrogen to ensure the jars and balls are submerged. Allow the jars and balls to cool until the rapid boiling of liquid nitrogen stops.
2. Using a large metal tongs, remove a single grinding jar and ball. Pour excess liquid nitrogen if present in the jars back into the styrofoam box.
3. Quickly add frozen cell pellet/PLB mixture and a grinding ball to the grinding jar and seal the jar.
4. Re-chill the jar in liquid nitrogen until boiling stops.
5. Loosen the grinding jar ¼ turn, place in a mixer mill and mill for 3 minutes at 15 Hz. Loosening the jar ¼ will allow any liquid nitrogen and gases escape from the jar during the 3-minute milling process.
6. Tighten the jar and return to liquid nitrogen.
7. Repeat for 5 more cycles.
8. Remove the jars from the mixer mill and chill the tips of two spatulas in liquid nitrogen.
9. Partly fill a 50 mL falcon tube with liquid nitrogen and place it upright in a liquid nitrogen bath. Open the grinding jar and recover the lysate powder into a 50 mL falcon tube using liquid nitrogen chilled spatulas, using a separate chilled spatula for each sample to avoid contamination.
10. Pierce the cap of the tube containing yeast lysate and place it upright in a -80°C freezer, until the liquid nitrogen evaporates. Samples can be stored at -80°C for extended periods.

#### 1.4 Lysate Clarification

- It is critical here to use a clean workspace, filter tips, with RNase-free gloves and tubes. Do this for all steps when working with RNA.
- Always keep RNA samples on ice between steps.

1. Thaw the yeast lysate powder gently (on ice or in a cold room) and transfer a 2 mL tube. Typical recovered yield should be ∼1.5 mL.
2. Immediately centrifuge at 3,000 x g, 4°C for 5 minutes.
3. Recover supernatant into 2 mL RNase-free tube.
4. Further clarify the supernatant by spinning at 20,000 x g, 4°C for 10 minutes and recover the supernatant again in a 1.5 mL tube.

#### 1.5 Lysate RNA quantification

- To quantify RNA concentration, use a Qubit fluorometer with BR (Broad-Range) assay kit or other.

1. Mix 199 µL RNA BR buffer and 1 µL BR reagent for each sample in a 1.5 mL tube.
2. Transfer 199 uL solution to Qubit assay tube,
3. Add 1 µL sample to the Qubit assay tube with 199 µL solution.
4. Briefly vortex and spin down on a microfuge.
5. Incubate at room temperature for 2 minutes.
6. Take readings on Qubit (select correct assay: RNA-BR (Broad Range)).

#### 1.6 Aliquot Lysates

- For ribosome profiling, multiple aliquots of 30 µg of total RNA into 1.5 mL tubes can be made and stored in -80°C. This leaves multiple backup samples should something go wrong.
- Additionally, aliquoted lysates provide extra material which can be used for RNA-seq to measure relative mRNA abundance.
- For an example of typical RNA yields from an experiment with 150 mL cultures (2% glucose minimal medium) of *K. marxianus* grown between A_600_ 0.6 - 1.0 see below.
- Each 150 mL culture should generally provide ∼500-900 µg RNA.

|  | concentration<br>(µg/ mL) | Volume (µL)<br>for 30 µg | polysome<br>buffer (µL) | total RNA yield (µg)<br>from 1.5 mL lysate |
| --- | --- | --- | --- | --- |
| sample 1 | 557 | 54 | 146 | 835 |
| sample 2 | 338 | 89 | 111 | 507 |
| sample 3 | 338 | 89 | 111 | 507 |
| sample 4 | 598 | 52 | 148 | 897 |
| sample 5 | 571 | 53 | 147 | 856 |
| sample 6 | 455 | 66 | 134 | 682 |

1. After RNA quantification, calculate the volume of clarified lysate required to aliquot 30 µg. In addition, note the volume of each sample.
2. Once 30 µg material has been dispensed into a tube, immediately drop it in a dewar flask of liquid nitrogen.
3. To collect the tubes, one may use a styrofoam box and a plastic sieve to quickly collect all tubes and place them into a freezer box.
4. Store the lysate stocks at -80°C indefinitely.

#### 2.1 Preparation of sucrose gradients

- Sucrose gradients for monosome isolation can be formed via two methods:

○ Using a Gradient master (Gradient Master 108, Biocomp) to quickly form reproducible gradients.
○ Bench-top gradient formation if a Gradient master is not available; however, this takes 4 hours for gradients to form.
- Alternatively, ribosomes can be pelleted in 1 M sucrose cushion as described in the Ingolia Protocol 2017 (McGlincy and Ingolia, 2017).

Gradient master method (preferred)

1. Prepare 10% and 50% sucrose solutions as described in the recipe section.
2. Place a polypropylene tube in the gradient master SW41ti metal marker supplied with the gradient master and using a fine-tip permanent marker, mark a line around the polypropylene tube.
3. Using a 50 mL syringe, carefully dispense 10% sucrose solution until the solution is raised to be in line with the marker.
4. Using a 10 mL syringe, carefully dispense 50% sucrose solution under the 10% sucrose and dispense until the 10%-50% sucrose border is lined up with the marker.
5. Carefully place the caps on each tube, using a syringe or P1000, carefully remove excess sucrose solution from the cap.
6. Turn on the gradient master and level the circular platform using the DOWN and UP options. When level, select DONE.
7. Select LIST, then SW41, navigate to “Short Sucr 10-50% wv 14S” and select USE and then RUN to begin gradient formation.
8. Place the tubes in a cold room for 45 minutes to allow gradients to cool before loading samples.

Bench-top method (alternative)

1. Prepare 10% and 50% sucrose solutions as described in solutions/buffers section.
2. Dispense 5.5 mL 50% sucrose solution to each polypropylene tube.
3. Gently layer 5.5 mL 10% sucrose on top of the 50% sucrose solution.
4. Seal the tubes with parafilm and gently lay the tubes horizontally for 4 hours.
5. Place the tubes vertically in the cold room for 45 minutes to allow gradients to cool before loading.

#### 2.2 RNase Digestion

1. Thaw 30 µg RNA samples on ice.
2. Dilute each 30 µg sample to 200 µL with ice-cold PB (polysome buffer, see solutions/buffers section).
3. To each sample, add 1.5 µL (10 U/µL) RNase I and incubate for 45 minutes at room temperature with agitation (T-shaker).
4. Place the samples on ice and proceed directly with ultracentrifugation.

#### 2.3 Ultracentrifugation

- For this protocol, use sucrose gradients and fractionation to isolate the monosome which contains the ribosome/RPF.
- Alternatively, if the above method is not available, have also pelleted monosomes using smaller scale ultracentrifugation. This alternative method is also described below.

Preferred method: Sucrose Gradient Ultracentrifugation

1. Turn on ultracentrifuge and input the following settings:

- Rotor: SW41-Ti
- Temperature: 4°C
- Speed: 36,000 RPM
- Time: 3 hours (3:00)
- Acceleration: 5
- Deceleration: 5
2. Pre-cool the ultracentrifuge by selecting the pre-cool option, the swinging bucket rotor and tubes are stored in a cold room.
3. In a cold room, remove 200 µL from the top of each gradient (200 µL is the volume to be added) and gently layer 200 µL of the sample using a P200.
4. Balance each gradient tube within 20 mg (0.02 g).
5. Place rotor in the ultracentrifuge and slide the door to close.
6. Press ENTER followed by START to begin the run.
7. Once the run is completed, allow air back into the rotor chamber by pressing Vacuum.
8. Once the pressurization is complete, slide open the centrifuge door and carefully remove the rotor and place it back on the rotor stand.
9. Carefully remove each tube and place on the provided rack.
10. Tubes can be left in the cold room or a fridge during the fractionation step.

Alternative method: Ribosome Pelleting Ultracentrifugation

1. Pre-cool the ultracentrifuge.
2. Pre-cool the TLA120.1 rotor in a cold room or fridge.
3. To each tube, pipette 200 µL of each sample.
4. Weigh and balance each tube with 10 mg.
5. Run at 120,000 rpm for 2 hours at 4°C.
6. Using a clean metal tongs, carefully remove each tube and using a P200, carefully remove the supernatant. Use a P10 to remove small volumes of remaining supernatant. A glassy pellet should be visible.
7. With this method, resuspend the pellet in 250 µL ultra-pure water and proceed directly to RNA extraction via Trizol protocol according to manufacturer’s instructions.

#### 2.4 Density Gradient Fractionation and Monosome Collection

- In this protocol, a tracerDaq is connected to a Windows desktop and Brandel UV detector, which provides a digital output of absorbance readings. Alternatively, the Brandel UV detector has a paper output which can be used as an alternative.

1. Assemble the fractionator equipment and prewarm the UV lamp, the lamp will be ready when the red light turns green.
2. Select the following settings on the pump and absorbance detector:

a. 1.5 mL/min flow speed
b. 0.2 Sensitivity
c. Baseline speed of 60 (if using paper graph output)
3. Using the glass syringe provided with the fractionator, uptake 60% CsCl solution and insert into the pump.
4. Once the lamp indicator is green, fill a polypropylene tube with Ultra-pure water, insert tube into the detector component and pierce the tube.
5. Pierce the polypropylene tube by raising the needle just below the polypropylene tube, then use the screw to raise the needle and pierce the tube.
6. Select normal & forward to begin pumping CsCl into the tube. The CsCl will raise the tube contents through the detector unit and out through the outflow tube. For this tube filled with ultra-pure water, a flat “blank” line will eventually form.
7. Using rapid reverse mode on the pump, remove some CsCl back into the syringe. Leave a noticeable amount remaining in the polypropylene tube so that the needle is submerged in CsCl.
8. Lower the needle using the screw, then lowering the needle stand, remove the polypropylene tube and discard.
9. When the tube is removed, start the forward pump to remove any air bubbles until CsCl comes through, then stop. Samples are now ready to be processed.
10. Once liquid comes through the outflow tube, carefully insert it into the 1^st^ well (A1) of a 96 well plate, hold for 12 seconds and move to the next and so on.
11. Disassemble all parts and wash with distilled water and leave them on a layer of tissue to dry on the bench. For the inflow tube (thick), wash with water. For the thinner outflow tube, use a P1000 to wash through 1 mL pure ethanol, then 1 mL water, then 2 x 1 mL empty pipette to push through any remaining liquid.
12. From the 96 well plate, the monosome fraction is selected using a UV reading from a plate reader. With readings from 260 nm, select the well with peaks corresponding to 80S. See supplementary figure 2 for an example of a trace from the fractionation.
13. Transfer the well(s) containing the monosome fraction to an RNase-free 1.5 mL tube. Typically, this is 2-3 wells, each containing ∼300 µL.
14. Proceed with the Trizol protocol to isolate RNA with the manufacturer’s instructions, resuspending in 12 µL of ultrapure water.

**Figure S2.**
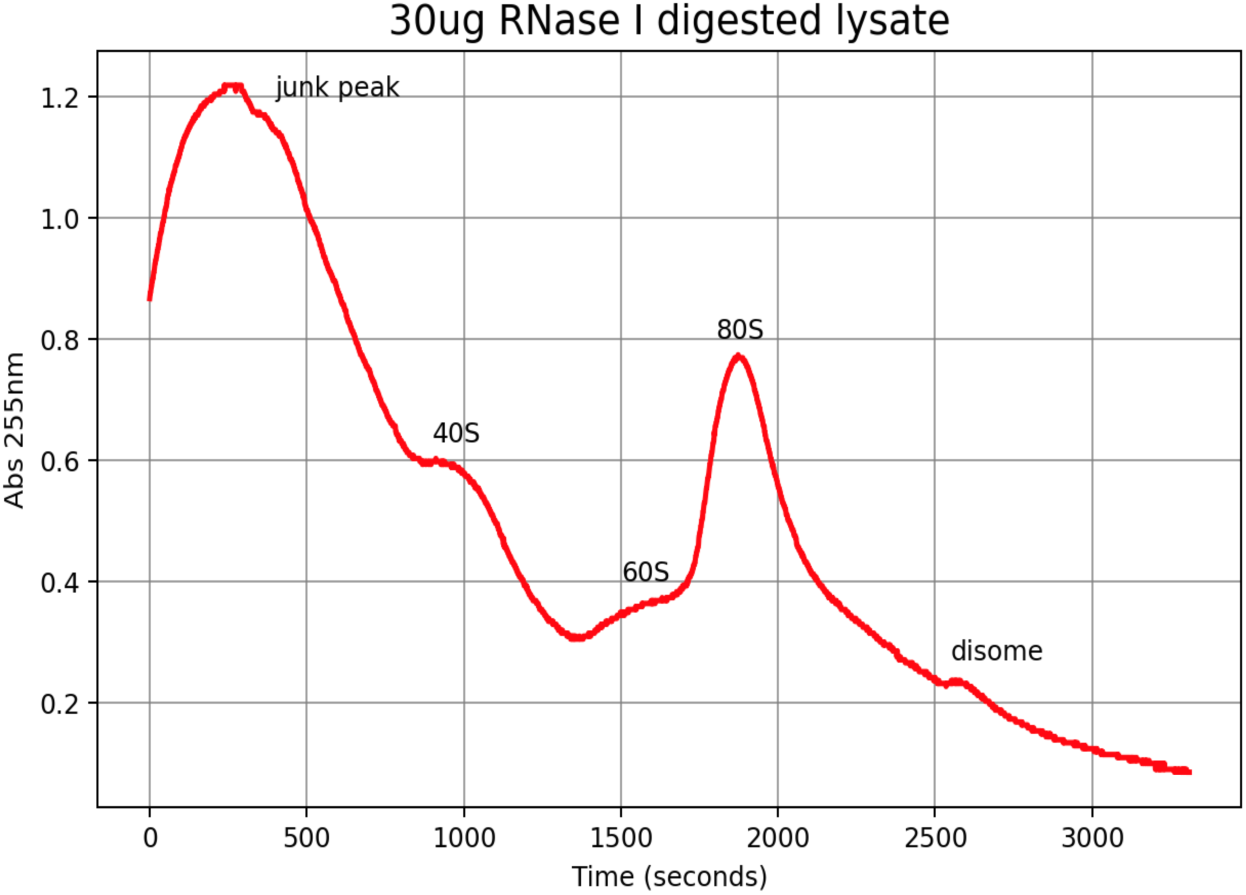
Sucrose gradient fractionation absorbance distribution of 30 µg RNase I digested material.

#### 2.5 RPF Size Selection (Urea-PAGE gel)

- After polysome RNase digestion and monosome isolation, the RPFs need to be size selected and purified. This is done with a size selection gel using a marker 26-34nt to select these range of fragments.
- In yeast, RPFs are ∼28nt in length and the use a 26-32nt marker allows excision of RPFs from a PAGE gel.
- For all gels required in this protocol, generally make a homemade gel mix described in the recipe table below. These stocks can be stored at room temperature for 2-3 months.
- Pre-cast gels are also available commercially which should be adequate for these experiments.
- Important note: The 28 and 32 nt RPF markers, which are used in high concentration, can contaminate libraries. It is recommended that careful dispensing of these markers is important to avoid contamination. In addition, allow empty wells between both marker and sample lanes to avoid cross contamination.

1. Prepare 15% Urea-PAGE gel stock solution as described in reagents. For a single gel, combine 7.478 mL Urea-PAGE stock solution (recipes), 18.75 µL APS and 3.75 µL TEMED to a 15 mL tube, vortex and using a 10 mL pipette, immediately add to 0.75 mm Bio-Rad gel plate sandwich and slowly insert 10 well comb. Leave to polymerize for 2 hours.
2. Set up the electrophoresis tank and add 1x TBE buffer.
3. Insert the polymerized gel and using a 10 mL syringe, clean the wells with buffer from the tank with 1X TBE 3 times. Be careful to not damage the gel separating the wells.
4. Pre-run at 300 V constant (∼15 mA) for 20-30 minutes (1x TBE)
5. Clean again with syringe.
6. Prepare marker: 1 µL marker + 7 µL water + 4 μL 3X RNA loading dye.
7. Prepare RPF sample: 12 μL RPF sample + 6 μL 3X RNA loading dye.
8. Denature marker and RPFs for 90 seconds at 80°C.
9. Load markers and RPF samples, leaving at least one lane between samples.
10. Run gel for 70 minutes at 300 V constant (∼15 mA).
11. Carefully remove gel and place into a tank with 20 mL 1X TBE and 2 μL SYBR Gold.
12. Place on a nutator for 2 minutes.
13. Visualize gel with Bio Rad GelDoc or equivalent. Figure 3 shows a typical gel from this experiment.

**Figure S3.**
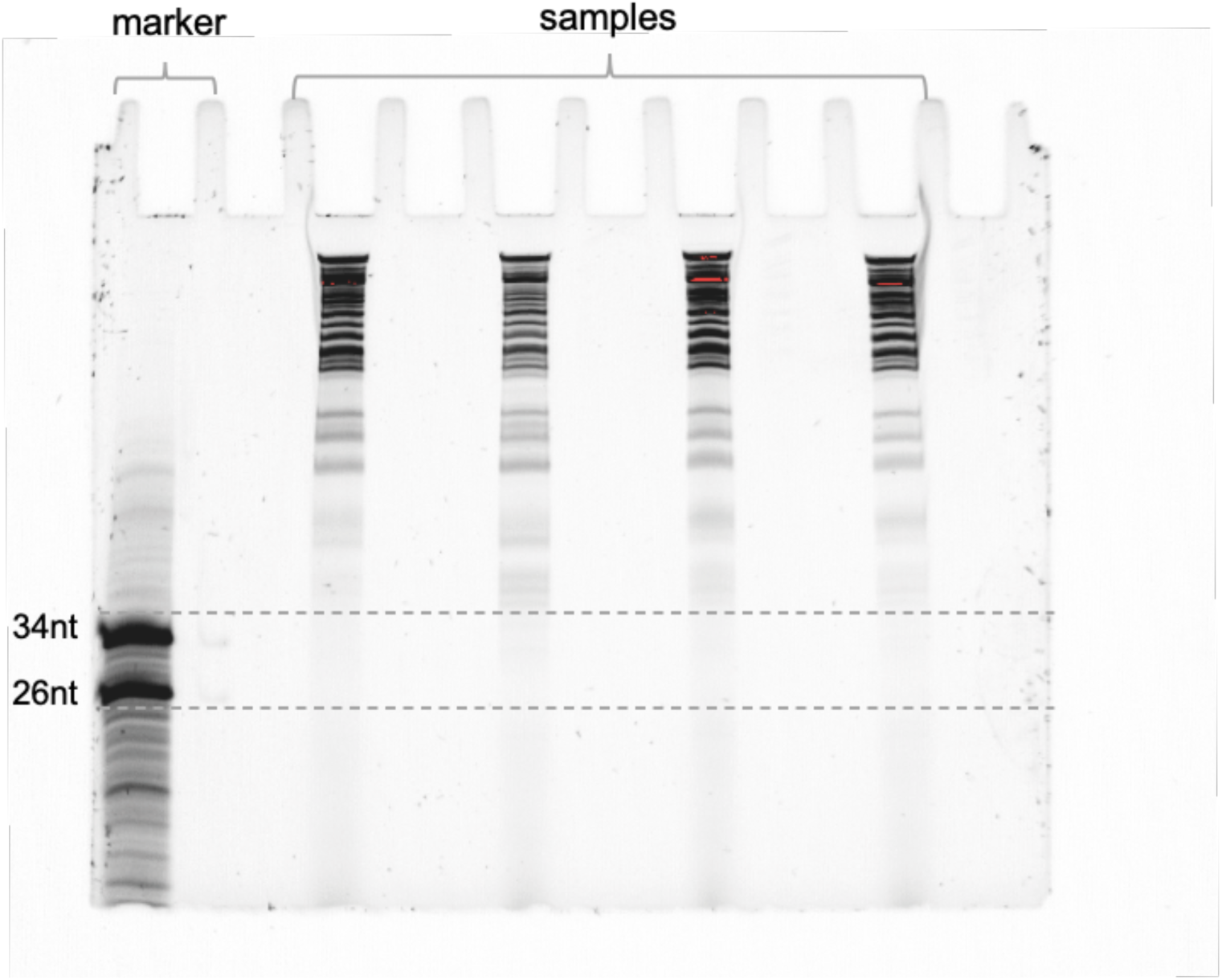
15% Urea-PAGE gel electrophoresis for RPF size selection with maker (left) and four loaded RNase I digested samples. Horizontal dashed gray lines represent the area for size selection across four footprint samples.

#### 2.6 RPF gel extraction

1. Cut out required gel section representing the RPFs. This can be done by printing an actual size image of the gel and marking the regions to be excised. The gel is then placed on a glass plate over the printed image and a scalpel is used to excise the region of interest. Alternatively, the gel can be placed under blue LED light and sliced using RPF markers as a guide.
2. Place each gel slice in a RNase-free 1.5 mL tube.
3. Add 500 µL RNA elution buffer.
4. Shake overnight on shaker at room temperature to extract RNA from slices at 1000 rpm.
5. Briefly centrifuge and collect the liquid, transfer to a new RNase-free 1.5 mL tube.
6. Add 500 µL of ice-cold isopropanol
7. Add 1.5 µL glycoblue.
8. Briefly vortex.
9. Precipitate in -20°C overnight or -80°C for 1 hour.
10. Thaw on ice and pellet by centrifugation for 30 minutes at 20,000g and 4°C.
11. Remove supernatant and wash pellet in 1 mL cold 80% ethanol.
12. Centrifuge again.
13. Carefully pipette all liquid from the tube and place it sideways on a rack in a chemical hood. Leave for 5 minutes to air dry remaining ethanol.
14. Resuspend in 4 µL ultra-pure water.
15. Store at -20°C overnight or -80°C indefinitely.

#### 3.1 Enzymatic pre-adenylation of linker using Mth RNA ligase

- Linker oligonucleotides requires enzymatic pre-adenylation prior to ligation to RNA fragments with T4 Rnl2(tr) K227Q, as this enzyme can only ligate adenylated linkers.

1. In a PCR tube, add the following:

a. 1.2μL linker oligonucleotide at 100 μM
b. 2μL 10X 5’ DNA adenylation action buffer
c. 2μL 1 mM ATP
d. 12.8μL water
e. 2μL Mth RNA Ligase
2. Incubate for 1 hour at 65°C, then heat-inactivate the enzyme by incubation at 85°C for 5 minutes.
3. Add 30 μL water to the sample and then purify using the Oligo Clean & Concentrator kit according to the manufacturer’s instructions, except elute in 6 μL nuclease-free water.
4. Store at −20°C and avoid repeated freeze–thaw.

#### 3.2 Dephosphorylation and Linker Ligation

- The purpose of dephosphorylation with T4 PNK is to “heal” the 3’ ends of RPFs as nuclease digestion introduces a 3’ cyclic phosphate which is incompatible for 3’ ligation to the pre-adenylated barcoded linker.
- As a positive control for dephosphorylation and linker ligation, use the 26 and 34 nucleotide RNA markers. These markers (like RPFs) require dephosphorylation prior to linker ligation and are thus suitable controls. See Figure 4B for positive control gel.
- To aid understanding of how this library preparation works, a molecular schematic of library preparation is provided in Figure 1.
- To avoid size-selection gel for ligation products, McGlincy and Ingolia 2017 use yeast 5’ deadenylase and RecJ exonuclease to digest un-ligated linkers from the sample, meaning no separation of ligated products from un-ligated-linkers via size selection is required. However, in this protocol, a size-selection gel is preferred as it allows monitoring of efficiency of the ligations.
- An example of a size selection gel to separate un-ligated linkers from ligated RPFs is provided in Figure 4A.

1. Set up each phosphorylation reaction as follows (for positive controls with RPF markers, substitute RNA sample with 0.5 μL RPF markers (100 μM) and 3 μL ultra-pure water):

| Component | Volume ($\mu\text{L}$) | Final |
| --- | --- | --- |
| RNA sample | 3.5 | NA |
| T4 PNK buffer (10X) | 0.5 | 1X |
| T4 PNK (10 U/$\mu\text{L}$) | 0.5 | 5 U |
| SUPERase•In | 0.5 | 10 U |
| Total volume | 5 |  |

2. Incubate for 1 hour at 37°C

3. Prepare the linker ligation reaction by adding components as follows directly to the dephosphorylation reaction, bringing the total volume to 10 μL.

| Component | Volume ($\mu\text{L}$) | Final |
| --- | --- | --- |
| Dephosphorylated RNA | 5 |  |
| 50% w/v PEG-8000 | 3.5 | 17.5% |
| 10X T4 RNA ligase buffer | 0.5 | 1X |
| Pre-adenylated linker (20 $\mu\text{M}$) | 0.5 | 1 $\mu\text{M}$ |
| T4 Rnl2(tr) K227Q (200 U/$\mu\text{L}$) | 0.5 | 100 U |
| Total volume | 10 |  |

4. Incubate the ligation reaction for 3 hours at 22°C (RT).

5. Run a 15% PAGE gel and size select for ligated products by following the previous RPF size selection gel and overnight extraction protocol, eluting ligated products with 11 μL ultra-pure water. In addition to ligation samples, load the ssRNA low range ladder (NEB) as a guide for size selection. Ligated products will be present as a band between the 50-80 nt ssRNA markers.

#### 3.3 Pooling and purification of ligations

- At this point, each individual sample contains a unique barcode within its linker. This allows for pooling of up to 8 individual samples with separate barcodes. This is advantageous as it reduces sample to sample variability during cDNA library preparation. Other advantages include increased material allows product bands to be more easily visualized on polyacrylamide gels.
- If doing a gel purification step of ligations, you may pool individual samples before loading onto gel.

1. Each sample to be pooled contributes 11 μL volume. Calculate the total sample volume: #samples x 11 μL.
2. If sample is <50 μL, bring total volume to 50 μL with ultra-pure water.
3. To the sample, add twice the volume of oligo binding buffer.
4. Add ethanol equal to 8 times the original total sample volume. eg, six samples (66 μL) use 528 μL ethanol.
5. Load samples onto the Zymo spin-column nested in a collection tube. Load no more than 800 μL at once (8 samples max).
6. Centrifuge the column for 30 seconds at 12,000 x g and discard the flow-through.
7. Add 750 μL DNA wash buffer and spin at 12,000 x g for 30 seconds.
8. Centrifuge again (with no wash) for 1min at max speed to remove any residual wash buffer.
9. Transfer the Zymo-Spin column into a 1.5 mL RNase-free tube and add 10 μL water.
10. Centrifuge for 30 seconds at 12,000 x g and recover RNA in the eluted liquid.
11. Store overnight at -20°C or indefinitely at -80°C.

#### 3.4 Size selection of ligation products

- The ligated RPFs are separated from the linker and non-ligated RNA fragments using a size selection gel.
- An example of PAGE separation of ligation products is visualized in Figure 4A, including RPF marker ligation in Figure 4B.
- RPFs (∼28 nt) ligated to 33 nt linker produces a ligated product ∼61 nt in size.

1. Carry out excision of ligation products as described previously for RPF size selection.
2. Isolate RPFs from gel slices as described previously, except resuspend RNA in 10 μL ultra-pure water.

**Figure S4.**
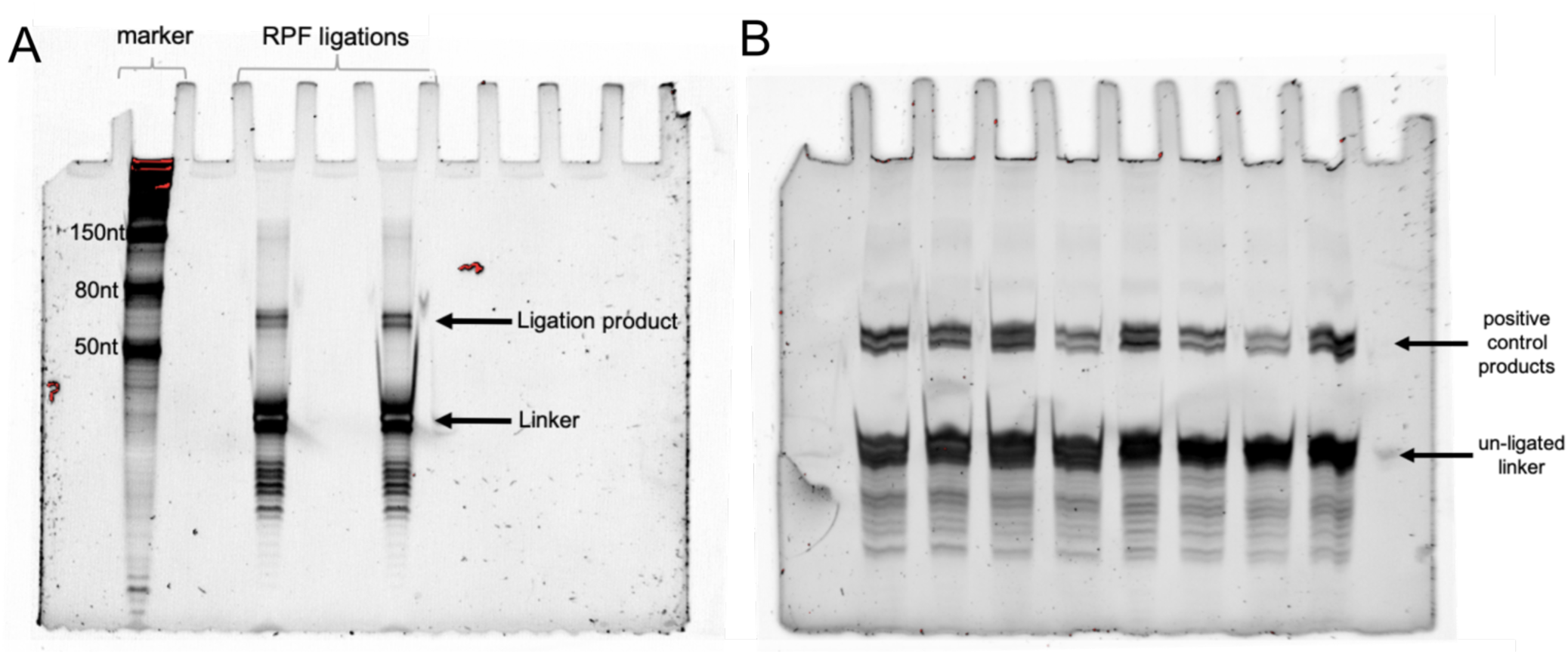
Polyacrylamide gel electrophoresis separation of ligation products. A. PAGE-gel separation of RPF samples with ssRNA marker. B. PAGE gel separation of positive control ligation products.

### 3.5 Reverse Transcription

- As a marker here, use the NEB Low Range ssRNA ladder.
- The expected product size is ∼111 nt.
- Temperatures have been optimized as described in McGlincy and Ingolia 2017 to reduce non-templated addition of nucleotides to cDNA products.

1. Add 2 μL reverse transcription primer (NI-802) at 1.25 μM to all RNA samples, bringing sample volume to 12 μL.
2. Denature for 5 min at 65°C in a PCR machine and then place on ice.
3. Cool the PCR machine to 50°C.
4. Set up the following reaction (if using Superscript, replace 1 μL Protoscript II with 1 μL Superscript III 200U/μL):

| Component | Volume ($\mu\text{L}$) | Final |
| --- | --- | --- |
| RNA sample and primer | 12 | NA |
| 5X Protoscript II buffer | 4 | 1X |
| dNTPs | 1 | 1 |
| 0.1M DTT | 1 | 5 mM |
| SUPERase•In (20U/ $\mu\text{L}$) | 1 | 20 U |
| Protoscript II (200U/ $\mu\text{L}$) | 1 | 20 U |

5. Incubate for 30min at 50°C (Protoscript II) or 55°C (Superscript III).
6. Hydrolyze the RNA template by adding 2.2 μL 1M NaOH to each tube and incubate at 70°C for 20 minutes.
7. Add 28 μL water, bringing total volume to 50 μL.
8. Purify sample using oligo clean and concentrator kit, except elute in 8 μL water.
9. Proceed to RT gel for size selection of RT products.

#### 3.6 Size selection of RT products

- Similar to previous PAGE gels, but 7.5%, not 15%.
- This gel separates the un-extended primer from the RT product see Figure 5 for example.

1. Prepare 7.5% PAGE gel as described in solutions/reagents.
2. Once polymerized, pre-run at 15mA (300V) for 20mins in 1X TBE.
3. Denature samples 90 seconds at 80°C.
4. Each sample: 8 μL cDNA + 4 μL 3X loading dye.
5. Use NEB ssRNA low range ladder as marker.
6. Run gel for 40 minutes.
7. Stain as previously described and visualize on GelDoc.
8. Cut out slice representing cDNA products and elute in 750 μL of DNA extraction buffer.
9. Place on shaker overnight at RT.
10. Transfer elution to 1.5 mL tube.
11. Add one volume of isopropanol and 1.5uL glycoblue.
12. Vortex briefly.
13. Precipitate for one hour in -80°C and resuspend in 12 μL ultra-pure water.
14. Thaw on ice and centrifuge at 20,000 x g for 20 minutes.
15. Discard supernatant and add 1 mL of ice-cold 80% ethanol.
16. Centrifuge again and discard supernatant.
17. Resuspend in 12 μL ultra-pure water.
18. Store cDNA products at -20°C or proceed directly to circularization reaction.

**Figure S5.**
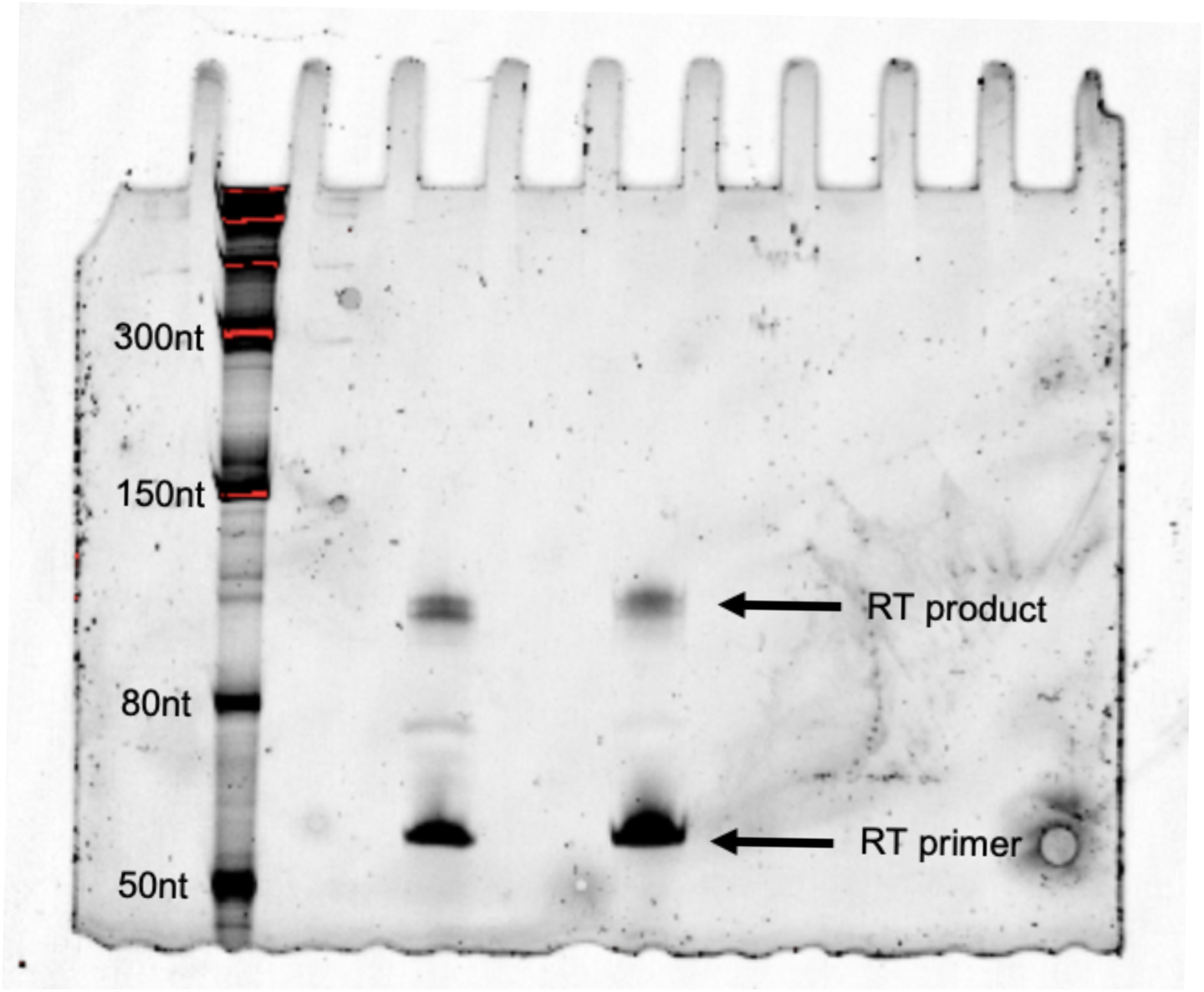
7.5% Urea-PAGE gel separation of reverse transcription product and primer. NEB ssRNA ladder is used as ladder.

#### 3.7 cDNA Circularization

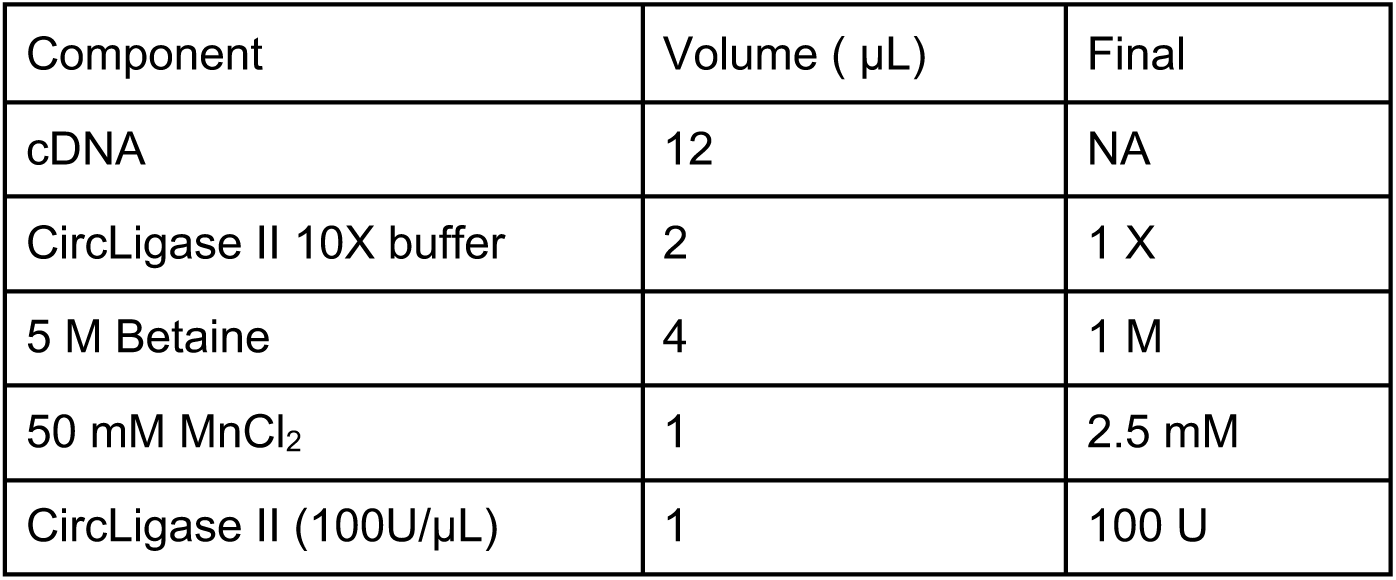

1. Transfer 12 μL RT product to a PCR tube.
2. Prepare circularization reactions as shown in table.
3. Incubate for 1 hour at 60°C in a thermal cycler.
4. Heat inactivates the enzyme at 80°C for 10 minutes.
5. The circularization products may be stored at -20°C.
6. At this point, proceed to rRNA removal via subtractive hybridization.

#### 3.8 rRNA depletion

- Table below describes rRNA depletion oligos. These are DNA oligos which are biotinylated at the 5’end and purified via HPLC.

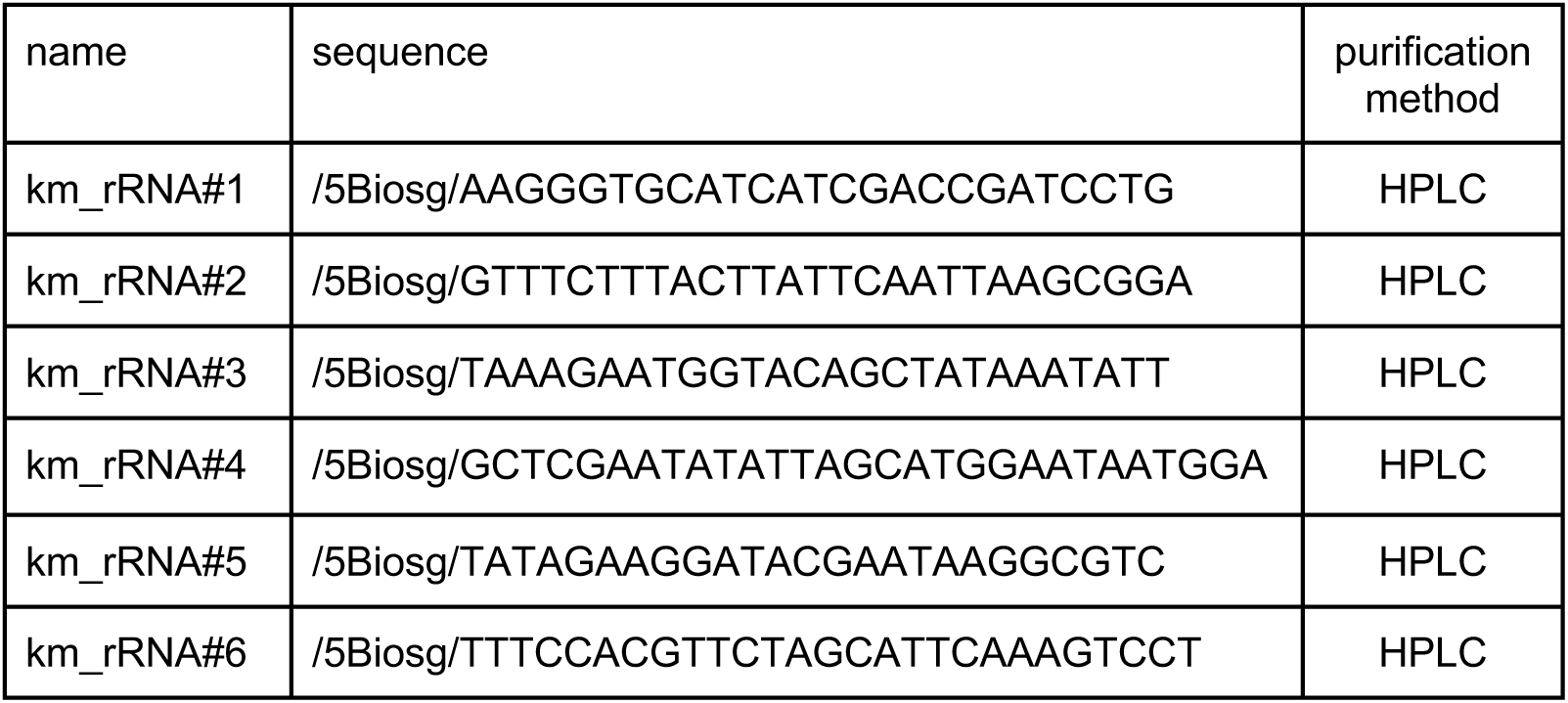

1. To make 100 µL of subtractive oligo mix:

- Prepare a 100 µM stock of each of the 6 oligos. Store at -20°C.
- Add 10 µL of each oligo to a tube with 40 µL water.
- Final concentration for oligos will be 60 µM, with 10 µM of each oligo.
2. Combine in a PCR tube:

a. 5µL circularization reaction (use 5 µL of previously prepared 20 µL circularization reaction)
b. 1µL subtractive oligo mix
c. 1µL 20X SSC
d. 3µL water in a PCR tube
3. Place the PCR tube in a thermal cycler and denature for 90 seconds at 100°C. Then anneal at by reducing the temperature by 0.1°C s^-1^ to 37°C (i.e., reduce to 37°C by 0.1°C per second). Incubate for 15 minutes at 37°C.
4. Vortex the streptavidin C1 Dynabeads to resuspend beads in storage solution.
5. Transfer the total required volume of beads to a clean tube in a magnetic stand: use 25 µL beads per reaction, plus an additional 12.5 µL beads (37.5 µL for single depletion reaction).
6. Leave for one minute and carefully aspirate all the liquid.
7. Remove from rack and add 1 volume 1X bind/wash buffer.
8. Repeat twice.
9. Place the beads on a magnetic rack for 1 minute to isolate beads and aspirate the final wash solution.
10. Resuspend in 0.4 volumes of 2X bind/wash buffer (15 µL for single sample).
11. Transfer one 10 µL aliquot of beads per subtraction reaction into another tube.
12. Place bead aliquots in the T-shaker at 37°C and equilibrate for 15 minutes.
13. Transfer 10 µL of subtraction reaction directly from the PCR tube in the thermal cycler (step 2) to a bead aliquot in the T-shaker. Incubate for 15 minutes at 37°C with mixing at 1,000 rpm.
14. Place tubes on a magnetic rack and isolate beads for 1 minute. Recover 17.5 µL eluate from the depletion and transfer to the new tube.
15. Add 1.5 µL glycoblue, 6 µL of 5 M NaCl and 74 µL water, followed by 150 µL isopropanol. Leave for 30 minutes on ice.
16. Centrifuge at 20,000 x g for 30 mins.
17. Remove supernatant and wash pellet with 1 mL ice-cold 80% ethanol.
18. Centrifuge and remove supernatant.
19. Air dry for a few minutes in chemical hood until residual ethanol evaporates.
20. Resuspend in 10 µL water.

#### 3.8 Final library PCRs

- For the final library PCR, Phusion high fidelity polymerase is used.
- The aim of initial trial PCRs is to determine the optimal cycle number, that is, the lowest PCR cycle to reduce PCR duplicates.
- For each PCR, it is important to have a particular ratio of template to volume, ideally no more than 5%. This strategy ensures that the primer concentration remains greater than 10 times the concentration of extended PCR product, and thus that in later cycles primer annealing and extension predominates over re-annealing of the two template strands.
- Once the optimal cycle number has been determined (see Figure 6A for trial PCR gel), one can increase the scale of PCR reaction and size select the final library PCR products (see Figure 6B for optimum cycle PCR).
- The forward primer is NI-798. Reverse primers are NI-799, 822–826, each include a different barcode to demultiplex samples.

Set up trial PCR reactions as follows:

- For each sample, prepare a 10 μL PCR reaction with 0.5 μL circularized template.
- Trial cycles: 8, 10, 12, 14 (scale as needed).

| Component | 10 μL reaction | 50 μL reaction | Final concentration |
| --- | --- | --- | --- |
| 5X Phusion HF buffer | 2 | 10 | 1X |
| dNTPs 10 mM each | 0.2 | 1 | 0.2 mM each |
| 10μM Forward Primer (NI-798) | 0.5 | 2.5 | 500 nM |
| 10μM Reverse Primer | 0.5 | 2.5 | 500 nM |
| Circularized cDNA template | 0.5 | 2.5 | - |
| Ultra-pure water | 6.2 | 31 | - |
| Phusion polymerase (2U/μL) | 0.1 | 0.5 | 1 U |

| Stage | Temperature | Time and Cycles |  |
| --- | --- | --- | --- |
| Initial Denaturation | 98oC | 30 Seconds |  |
| Denaturation | 98oC | 10 Seconds | 7-14 cycles |
| Annealing | 65oC | 10 Seconds |  |
| Extension | 72oC | 5 Seconds |  |
| Final Extension | 72oC | 5 Minutes |  |
| Hold | 4oC | Until removed. |  |

The forward primer is NI-798. Reverse primers are NI-799, 822–826

Once the optimal desired PCR cycle has been determined, prepare a 50 μL reaction with 2.5 μL template.

#### 3.9 Final PCR gel

1. Prepare 8% PAGE-gel solution in Solutions/Buffers section.
2. Assemble gel tank and add 1X TBE.
3. Insert polymerized gel into tank and gel wells as described previously.
4. As a marker, use 1 μL of 100 bp ladder (dsDNA).
5. To a 50 μL PCR reaction, add 10 μL 6X DNA loading dye (for trial PCRs, add 2 μL of 6X DNA loading dye to 10 μL trial PCR).
6. Load gel by dividing the 50 μL reaction into 3 separate wells.
7. Run gel for 40 minutes at 180 V.
8. Stain gel for 3 minutes with 1X SYBR gold in 20 mL 1X TBE.

#### 3.10 Library DNA extraction

1. Cut product bands and continue with an overnight gel extraction by adding 750 μL DNA gel extraction buffer.
2. Leave on a shaker overnight.
3. Spin down tubes briefly and transfer the buffer to fresh 1.5 mL tube.
4. Add 1.5 μL glycoblue and 750 μL isopropanol. Vortex briefly.
5. Spin for 30 minutes at 20,000 x g for 30 minutes at 4°C and carefully discard supernatant.
6. Add 750 μL ice-cold 80% ethanol and spin again for 15 minutes and discard the supernatant.
7. Let the pellet air dry for 5 minutes in a vacuum hood.
8. Resuspend pellet in 15 μL ultra-pure water.
9. Store at -20°C.

Once products have been size selected and eluted, measure library concentration with Qubit (HS-dsDNA assay) or equivalent. ng/μL may be converted to nM using the following formula:

DNA concentration (nM) = (ng/μL)/(660 g/mol x library size) x 10^6^

**Figure S6.**
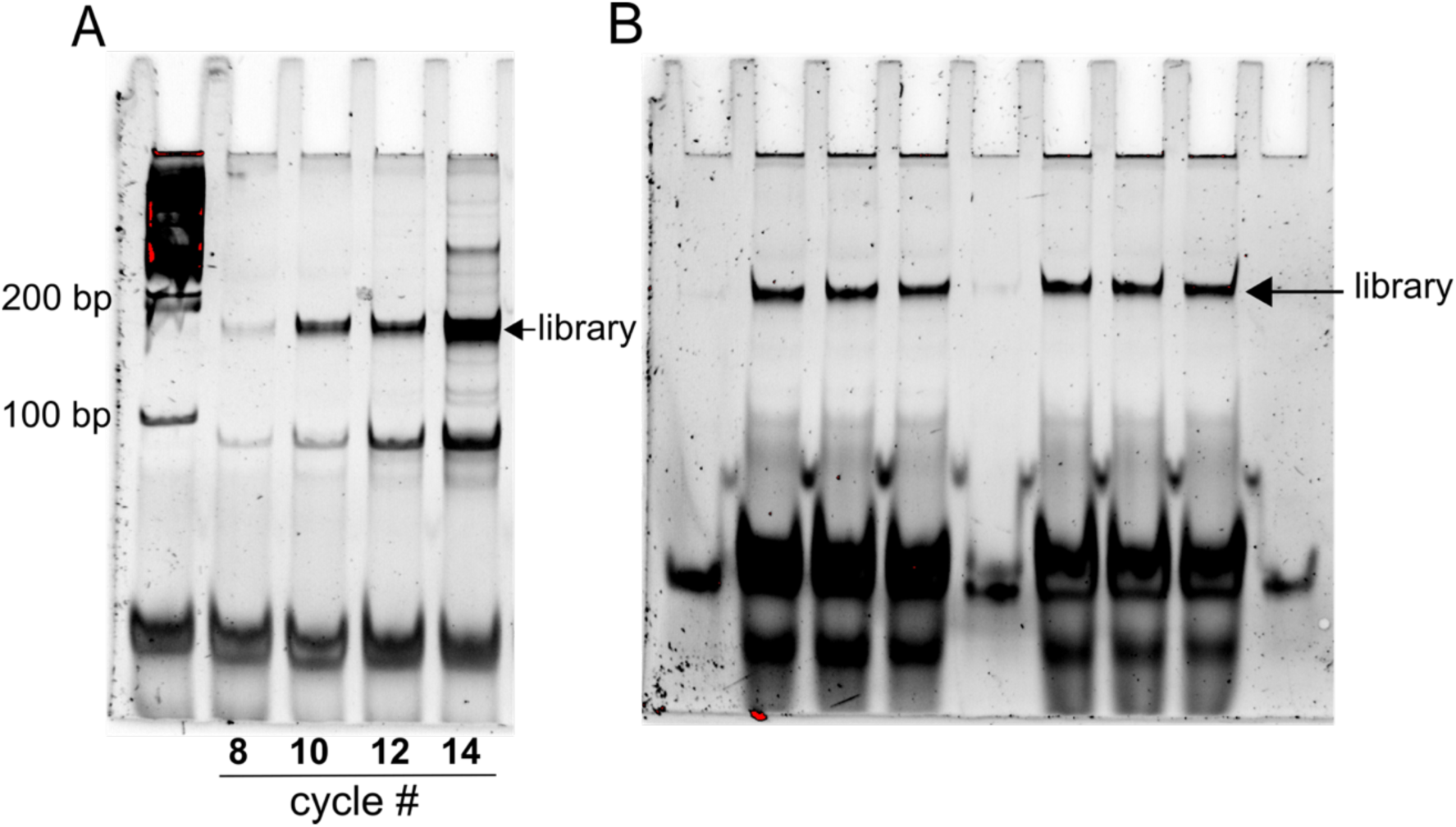
Final cDNA library PCR. A. Trial PCR with 8, 10, 12 and 14 cycles, leftmost lane contains 100 bp dsDNA ladder. 100 bp dsDNA used as ladder. B. PCR amplification with 8 cycles divided into 3 wells per sample.

#### 3.10 QC of Libraries

- Libraries for next generation sequencing are often loaded onto an Agilent Bioanalyzer to determine correct library size, prior to loading onto a flow cell.
- For our experiments, libraries are sequenced by the Genomics & Cell Characterization Core Facility (GC3F), University of Oregon on an Illumina HiSeq4000.
- Figure 7 shows an example of Agilent Bioanalyzer of library size distribution for a Ribo-Seq library.

**Figure S7.**
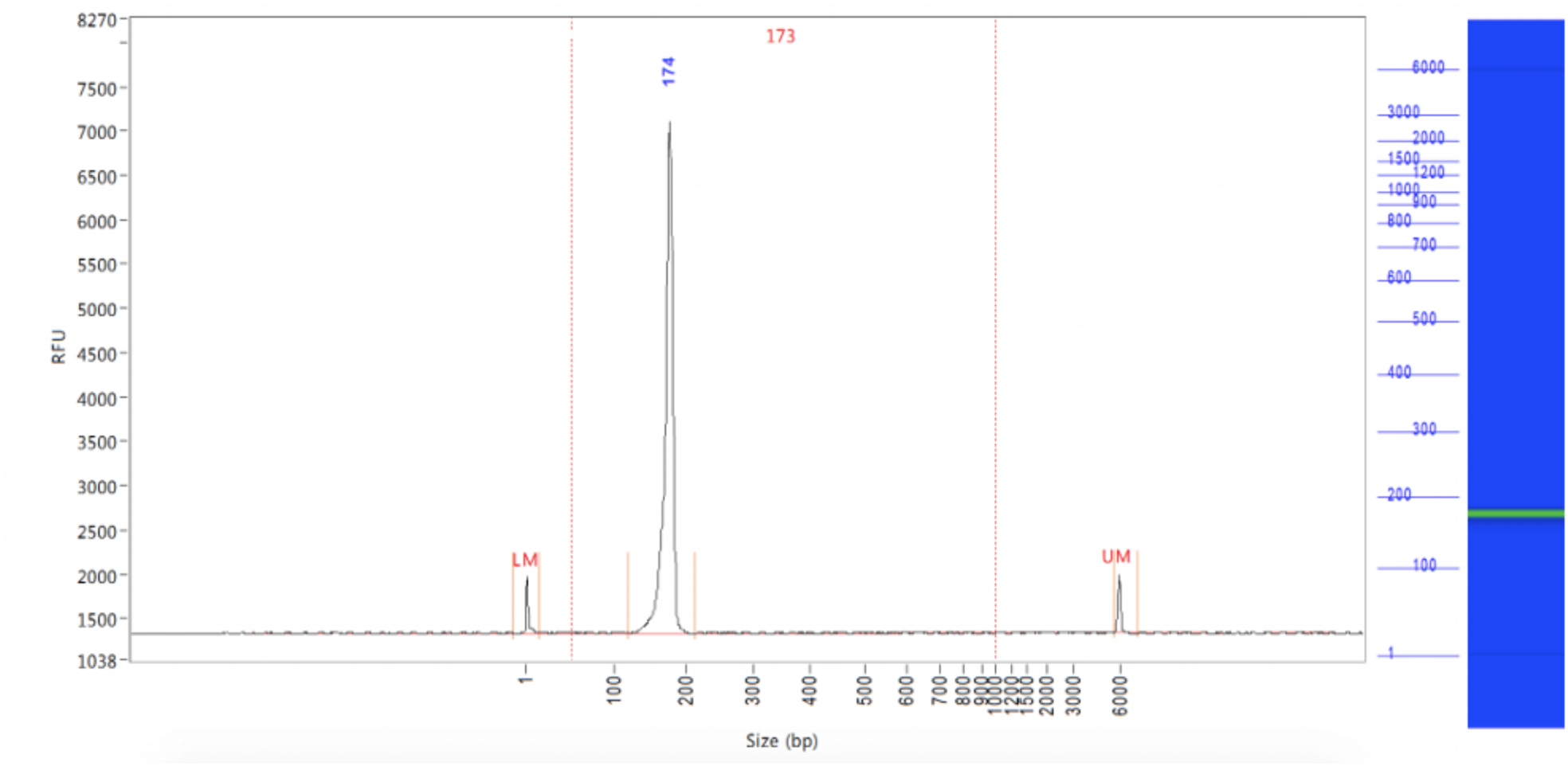
Bioanalyzer distribution of library size including lower and upper bioanalyzer markers. Ribosome profiling library is present at ∼174 size (bp).

## Bioinformatic Analysis

- For reference genome, *K. marxianus* DMKU3-1042 (Lertwattanasakul *et al*., 2015) is used. This includes the recently reannotated the DMKU3-1042 genome using ribosome profiling data which included over 150 novel genes and multiple gene annotation corrections (Bioarchive citation).
- All analysis of ribosome profiling and RNA-seq data is performed with Ubuntu server.
- The following software for processing the ribosome profiling data is provided before and usually installed via the conda environment package manager (https://conda.io/) (Anaconda software distribution, *Conda*).

○ FastQC to determine basic quality metrics such as read lengths, sequencing quality, GC content and more (Simon Andrews, Babraham institute).
○ Cutadapt is used to demultiplex and trim adapter sequences, specifically the Illumina adapters (Martin, 2011).
○ Bowtie is an aligner used to remove rRNA contaminants from the library followed by alignment to the *K. marxianus* strain DMKU3-1042 genome (Langmead *et al*., 2009).
○ Samtools converts the alignment SAM file to a sorted and indexed BAM alignment file (Li *et al*., 2009).
○ HT-seq counts the number of RPFs aligned to each gene using a GTF file (Anders, Pyl and Huber, 2015).

▪ From HT-SEQ, a count file is generated which can be used to determine sample correlations and for differential gene expression analysis.
○ bedtools coverts the BAM file to a forward and reverse strand genome alignment coverage file (Quinlan and Hall, 2010).
○ bedGraphToBigWig converts the coverage files to BigWig format which can be loaded onto a genome browser such as GWIPs-Viz (Michel *et al*., 2014).
○ For differential gene expression analysis, use DESeq2 (Love, Huber and Anders, 2014).
- Below, an example of commands used to process data via a Linux OS (Ubuntu) server is provided. Using these commands, one should be able to clip sequencing adapters, remove other rRNA contaminants and align sequences to the genome. A genome alignment is a precursor to generation of a counts table, which details the number of RPFs per gene for a specific condition, these tables are commonly used to determine reproducibility of replicates and for differential expression analysis. In addition, the genome alignment can be used to generate a bigWig file, which can be displayed as a private track on GWIPS-Viz.
- Figure 8 displays structure of a single read including UMIs, RPF, barcode and common adaptor. These reads are typically provided from a sequencing centre, which with clip Illumina adapters before providing the resulting reads to a customer.

Demultiplex and clip adapter, a fasta file describing barcodes is required (example below).

**Figure S8.**
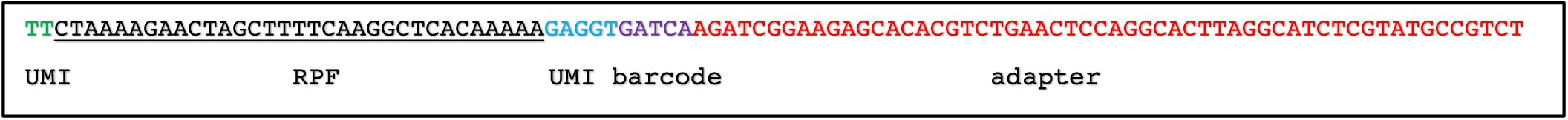
Structure of a single FASTQ sequence from HiSeq4000, generated with the ribosome profiling protocol described. During data processing, sequences upstream and downstream of the ribosome protected fragment “RPF” are clipped to allow alignment to the genome or transcriptome.

cutadapt -e 0.15 --no-indels -g file:barcodes.fasta -o “trimmed-{name}.fastq” test.fastq

>CTAGA_barcode

CTAGAAGATCGGAAGAGCACACGTCTGAA

>CGTAA_barcode

CGTAAAGATCGGAAGAGCACACGTCTGAA

Once demultiplexed, clip barcode and UMI.

cutadapt -u 2 -u -5 -m 25 -M 32 -j 12 -o example.fastq example.fastq

Remove remaining rRNA contaminants

bowtie -p 8 --un $1.temp path/to/rRNA/index $1 > non_coding.fastq

Align remaining reads to the genome

bowtie -p 2 -n 2 -m 1 -S path/to/genome/index example.fastq > genome_aligned.fastq

Convert sam to bam

samtools view -@ 32 -Sb example.sam > example.bam

Sort bam file

samtools sort -O bam example.bam > example_sorted.bam

Generate an index file

samtools index example_sorted.bam

Generate a counts table of mapped reads per gene (need for differential expression analysis)

htseq-count --format bam example_sorted.bam path/to/gtf > example_counts.tsv

Create coverage files. Here, split the forward and reverse strand reads. The chrominfo.tsv is a file describing chromosome names and length (example below).

bedtools genomecov -ibam $1_sorted.bam -strand + -g chromInfo.tsv -bg > example_FR.cov
bedtools genomecov -ibam $1_sorted.bam -strand - -g chromInfo.tsv -bg > example_RV.cov

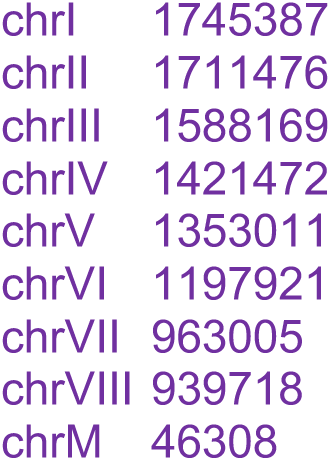

Create bigWigs for genome browser

bedGraphToBigWig example_FR.cov chromInfo.txt example_FR.bigWig
bedGraphToBigWig example_RV.cov chromInfo.txt example_RV.bigWig

## References

Alva TR, Riera M, Chartron JW. Translational landscape and protein biogenesis demands of the early secretory pathway in Komagataella phaffii. Microb Cell Fact 2021;20:19.

Andreev DE, O’Connor PBF, Loughran G et al. Insights into the mechanisms of eukaryotic translation gained with ribosome profiling. Nucleic Acids Res 2017;45:513–26.

Arevalo-Villena M, Briones-Perez A, Corbo MR et al. Biotechnological application of yeasts in food science: Starter cultures, probiotics and enzyme production. J Appl Microbiol 2017;123:1360–72.

Blevins WR, Tavella T, Moro SG et al. Extensive post-transcriptional buffering of gene expression in the response to severe oxidative stress in baker’s yeast. Sci Rep 2019;9:11005.

Brar GA, Weissman JS. Ribosome profiling reveals the what, when, where and how of protein synthesis. Nat Rev Mol Cell Biol 2015;16:651–64.

Broach JR. Nutritional Control of Growth and Development in Yeast. Genetics 2012;192:73 LP – 105.

Cernak P, Estrela R, Poddar S et al. Engineering Kluyveromyces marxianus as a Robust Synthetic Biology Platform Host. MBio 2018;9, DOI: 10.1128/mBio.01410-18.

Chomczynski P, Sacchi N. The single-step method of RNA isolation by acid guanidinium thiocyanate–phenol–chloroform extraction: twenty-something years on. Nat Protoc 2006;1:581–5.

Coloretti F, Chiavari C, Luise D et al. Detection and identification of yeasts in natural whey starter for Parmigiano Reggiano cheese-making. Int Dairy J 2017;66:13–7.

Delvigne F, Zune Q, Lara AR et al. Metabolic variability in bioprocessing: implications of microbial phenotypic heterogeneity. Trends Biotechnol 2014;32:608–16.

Doughty TW, Domenzain I, Millan-Oropeza A et al. Stress-induced expression is enriched for evolutionarily young genes in diverse budding yeasts. Nat Commun 2020;11:2144.

Duncan CDS, Mata J. The translational landscape of fission-yeast meiosis and sporulation. Nat Struct Mol Biol 2014;21:641–7.

Eisenberg AR, Higdon AL, Hollerer I et al. Translation Initiation Site Profiling Reveals Widespread Synthesis of Non-AUG-Initiated Protein Isoforms in Yeast. Cell Syst 2020;11:145–160.e5.

Fonseca GG, Heinzle E, Wittmann C et al. The yeast Kluyveromyces marxianus and its biotechnological potential. Appl Microbiol Biotechnol 2008;79:339–54.

Fu X, Li P, Zhang L et al. Understanding the stress responses of Kluyveromyces marxianus after an arrest during high-temperature ethanol fermentation based on integration of RNA-Seq and metabolite data. Appl Microbiol Biotechnol 2019;103:2715–29.

Gao J, Yuan W, Li Y et al. Transcriptional analysis of Kluyveromyces marxianus for ethanol production from inulin using consolidated bioprocessing technology. Biotechnol Biofuels 2015;8:115.

Gibson UE, Heid CA, Williams PM. A novel method for real time quantitative RT-PCR. Genome Res 1996;6:995–1001.

Groeneveld P, Stouthamer AH, Westerhoff H V. Super life--how and why “cell selection” leads to the fastest-growing eukaryote. FEBS J 2009;276:254–70.

Hahn S, Young ET. Transcriptional Regulation in *Saccharomyces cerevisiae*: Transcription Factor Regulation and Function, Mechanisms of Initiation, and Roles of Activators and Coactivators. Genetics 2011;189:705 LP – 736.

Hinnebusch AG. Translational regulation of GCN4 and the general amino acid control of yeast. Annu Rev Microbiol 2005;59:407–50.

Ingolia NT, Brar GA, Rouskin S et al. The ribosome profiling strategy for monitoring translation in vivo by deep sequencing of ribosome-protected mRNA fragments. Nat Protoc 2012;7:1534–50.

Ingolia NT, Ghaemmaghami S, Newman JRS et al. Genome-wide analysis in vivo of translation with nucleotide resolution using ribosome profiling. Science 2009;324:218– 23.

Ingolia NT, Hussmann JA, Weissman JS. Ribosome Profiling: Global Views of Translation. Cold Spring Harb Perspect Biol 2019;11, DOI: 10.1101/cshperspect.a032698.

Jiménez-Gutiérrez E, Alegría-Carrasco E, Sellers-Moya Á et al. Not just the wall: the other ways to turn the yeast CWI pathway on. Int Microbiol 2020;23:107–19.

Karim A, Gerliani N, Aïder M. Kluyveromyces marxianus: An emerging yeast cell factory for applications in food and biotechnology. Int J Food Microbiol 2020;333:108818.

Kiniry SJ, Judge CE, Michel AM et al. Trips-Viz: an environment for the analysis of public and user-generated ribosome profiling data. Nucleic Acids Res 2021;49:W662–70.

Kiniry SJ, O’Connor PBF, Michel AM et al. Trips-Viz: a transcriptome browser for exploring Ribo-Seq data. Nucleic Acids Res 2019;47:D847–52.

Kwon D-H, Park J-B, Hong E et al. Ethanol production from xylose is highly increased by the Kluyveromyces marxianus mutant 17694-DH1. Bioprocess Biosyst Eng 2019;42:63–70.

de la Torre-Ruiz MA, Pujol N, Sundaran V. Coping with oxidative stress. The yeast model. Curr Drug Targets 2015;16:2–12.

Lane MM, Morrissey JP. Kluyveromyces marxianus: A yeast emerging from its sister’s shadow. Fungal Biol Rev 2010;24:17–26.

Langmead B, Trapnell C, Pop M et al. Ultrafast and memory-efficient alignment of short DNA sequences to the human genome. Genome Biol 2009;10:R25.

Lertwattanasakul N, Kosaka T, Hosoyama A et al. Genetic basis of the highly efficient yeast Kluyveromyces marxianus: complete genome sequence and transcriptome analyses. Biotechnol Biofuels 2015;8:47.

Liu Y, Nielsen J. Recent trends in metabolic engineering of microbial chemical factories. Curr Opin Biotechnol 2019;60:188–97.

Ljungdahl PO, Daignan-Fornier B. Regulation of Amino Acid, Nucleotide, and Phosphate Metabolism in *Saccharomyces cerevisiae* Genetics 2012;190:885 LP – 929.

Marcet-Houben M, Gabaldón T. Beyond the Whole-Genome Duplication: Phylogenetic Evidence for an Ancient Interspecies Hybridization in the Baker’s Yeast Lineage. PLOS Biol 2015;13:e1002220.

Martin M. Cutadapt removes adapter sequences from high-throughput sequencing reads. EMBnet.journal 2011;17:10–2.

Martínez-Montañés F, Pascual-Ahuir A, Proft M. Toward a genomic view of the gene expression program regulated by osmostress in yeast. OMICS 2010;14:619–27.

Masser AE, Ciccarelli M, Andréasson C. Hsf1 on a leash – controlling the heat shock response by chaperone titration. Exp Cell Res 2020;396:112246.

McGlincy NJ, Ingolia NT. Transcriptome-wide measurement of translation by ribosome profiling. Methods 2017;126:112–29.

McManus CJ, May GE, Spealman P et al. Ribosome profiling reveals post-transcriptional buffering of divergent gene expression in yeast. Genome Res 2014;24:422–30.

Merrick WC. eIF4F: A Retrospective *. J Biol Chem 2015;290:24091–9.

Michel AM, Fox G, M Kiran A et al. GWIPS-viz: development of a ribo-seq genome browser. Nucleic Acids Res 2014;42:D859–64.

Mo W, Wang M, Zhan R et al. Kluyveromyces marxianus developing ethanol tolerance during adaptive evolution with significant improvements of multiple pathways. Biotechnol Biofuels 2019;12:63.

Mohammad F, Green R, Buskirk AR. A systematically-revised ribosome profiling method for bacteria reveals pauses at single-codon resolution. Elife 2019;8, DOI: 10.7554/eLife.42591.

Monteuuis G, Miścicka A, Świrski M, et al. Non-canonical translation initiation in yeast generates a cryptic pool of mitochondrial proteins. Nucleic Acids Res 2019;47:5777–91.

Morano KA, Grant CM, Moye-Rowley WS. The Response to Heat Shock and Oxidative Stress in *Saccharomyces cerevisiae* Genetics 2012;190:1157 LP – 1195.

de Nadal E, Posas F. Multilayered control of gene expression by stress-activated protein kinases. EMBO J 2010;29:4–13.

Nandy SK, Srivastava RK. A review on sustainable yeast biotechnological processes and applications. Microbiol Res 2018;207:83–90.

Parapouli M, Vasileiadis A, Afendra A-S, et al. Saccharomyces cerevisiae and its industrial applications. AIMS Microbiol 2020;6:1–31.

Rajkumar AS, Morrissey JP. Rational engineering of Kluyveromyces marxianus to create a chassis for the production of aromatic products. Microb Cell Fact 2020;19:207.

Rajkumar AS, Varela JA, Juergens H et al. Biological Parts for Kluyveromyces marxianus Synthetic Biology . Front Bioeng Biotechnol 2019;7:97.

Sanz AB, García R, Rodríguez-Peña JM et al. The CWI Pathway: Regulation of the Transcriptional Adaptive Response to Cell Wall Stress in Yeast. J fungi (Basel, Switzerland) 2017;4, DOI: 10.3390/jof4010001.

Schabort DTWP, Letebele PK, Steyn L, et al. Differential RNA-seq, Multi-Network Analysis and Metabolic Regulation Analysis of Kluyveromyces marxianus Reveals a Compartmentalised Response to Xylose. PLoS One 2016;11:e0156242.

Schena M, Shalon D, Davis RW et al. Quantitative monitoring of gene expression patterns with a complementary DNA microarray. Science 1995;270:467–70.

Sharma P, Wu J, Nilges BS et al. Humans and other commonly used model organisms are resistant to cycloheximide-mediated biases in ribosome profiling experiments. Nat Commun 2021;12:5094.

Smith JE, Alvarez-Dominguez JR, Kline N et al. Translation of small open reading frames within unannotated RNA transcripts in Saccharomyces cerevisiae. Cell Rep 2014;7:1858–66.

Spealman P, Naik AW, May GE et al. Conserved non-AUG uORFs revealed by a novel regression analysis of ribosome profiling data. Genome Res 2018;28:214–22.

Steitz JA. Polypeptide chain initiation: nucleotide sequences of the three ribosomal binding sites in bacteriophage R17 RNA. Nature 1969;224:957–64.

Sui Y, Wisniewski M, Droby S et al. Responses of yeast biocontrol agents to environmental stress. Appl Environ Microbiol 2015;81:2968–75.

Takors R. Scale-up of microbial processes: impacts, tools and open questions. J Biotechnol 2012;160:3–9.

Taymaz-Nikerel H, Cankorur-Cetinkaya A, Kirdar B. Genome-Wide Transcriptional Response of Saccharomyces cerevisiae to Stress-Induced Perturbations. Front Bioeng Biotechnol 2016;4:17.

Verduyn C, Postma E, Scheffers WA et al. Effect of benzoic acid on metabolic fluxes in yeasts: A continuous-culture study on the regulation of respiration and alcoholic fermentation. Yeast 1992;8:501–17.

Wang D, Wu D, Yang X et al. Transcriptomic analysis of thermotolerant yeast Kluyveromyces marxianus in multiple inhibitors tolerance. RSC Adv 2018;8:14177–92.

Wang Z, Gerstein M, Snyder M. RNA-Seq: a revolutionary tool for transcriptomics. Nat Rev Genet 2009;10:57–63.

Wehrs M, Tanjore D, Eng T et al. Engineering Robust Production Microbes for Large-Scale Cultivation. Trends Microbiol 2019;27:524–37.

Wolfe KH, Shields DC. Molecular evidence for an ancient duplication of the entire yeast genome. Nature 1997;387:708–13.

## References

Anders, S., Pyl, P. T. and Huber, W. (2015) ‘HTSeq--a Python framework to work with high-throughput sequencing data’, Bioinformatics (Oxford, England). 2014/09/25, 31(2), pp. 166–169. doi: 10.1093/bioinformatics/btu638.

Langmead, B. et al. (2009) ‘Ultrafast and memory-efficient alignment of short DNA sequences to the human genome’, Genome Biology, 10(3), p. R25. doi: 10.1186/gb-2009-10-3-r25.

Lertwattanasakul, N. et al. (2015) ‘Genetic basis of the highly efficient yeast Kluyveromyces marxianus: complete genome sequence and transcriptome analyses.’, Biotechnology for biofuels, 8, p. 47. doi: 10.1186/s13068-015-0227-x.

Li, H. et al. (2009) ‘The Sequence Alignment/Map format and SAMtools’, Bioinformatics (Oxford, England). 2009/06/08, 25(16), pp. 2078–2079. doi: 10.1093/bioinformatics/btp352.

Love, M. I., Huber, W. and Anders, S. (2014) ‘Moderated estimation of fold change and dispersion for RNA-seq data with DESeq2’, Genome Biology, 15(12), p. 550. doi: 10.1186/s13059-014-0550-8.

Martin, M. (2011) ‘Cutadapt removes adapter sequences from high-throughput sequencing reads’, EMBnet.journal, 17(1), pp. 10–12. doi: 10.14806/ej.17.1.200.

McGlincy, N. J. and Ingolia, N. T. (2017) ‘Transcriptome-wide measurement of translation by ribosome profiling.’, Methods (San Diego, Calif.), 126, pp. 112–129. doi: 10.1016/j.ymeth.2017.05.028.

Michel, A. M. et al. (2014) ‘GWIPS-viz: development of a ribo-seq genome browser.’, Nucleic acids research, 42(Database issue), pp. D859–64. doi: 10.1093/nar/gkt1035.

Quinlan, A. R. and Hall, I. M. (2010) ‘BEDTools: a flexible suite of utilities for comparing genomic features.’, Bioinformatics (Oxford, England), 26(6), pp. 841–842. doi: 10.1093/bioinformatics/btq033.

Verduyn, C. et al. (1992) ‘Effect of benzoic acid on metabolic fluxes in yeasts: A continuous-culture study on the regulation of respiration and alcoholic fermentation’, Yeast, 8(7), pp. 501–517. doi: 10.1002/yea.320080703.

